# The widespread influence of ZSWIM8 on microRNAs during mouse embryonic development

**DOI:** 10.1101/2023.06.21.545803

**Authors:** Charlie Y. Shi, Lara E. Elcavage, Raghu R. Chivukula, Joanna Stefano, Benjamin Kleaveland, David P. Bartel

## Abstract

MicroRNAs (miRNAs) pair to sites in mRNAs to direct the degradation of these RNA transcripts. Conversely, certain RNA transcripts can direct the degradation of particular miRNAs. This target-directed miRNA degradation (TDMD) requires the ZSWIM8 E3 ubiquitin ligase. Here, we report the function of ZSWIM8 in the mouse embryo. *Zswim8*^−/−^ embryos were smaller than their littermates and died near the time of birth. This highly penetrant perinatal lethality was apparently caused by a lung sacculation defect attributed to failed maturation of alveolar epithelial cells. Some mutant individuals also had heart ventricular septal defects. These developmental abnormalities were accompanied by aberrant accumulation of >50 miRNAs observed across 12 tissues, which often led to enhanced repression of their mRNA targets. These ZSWIM8-sensitive miRNAs were preferentially produced from genomic miRNA clusters, and in some cases, ZSWIM8 caused a switch in the dominant strand that accumulated from a miRNA hairpin—observations suggesting that TDMD provides a mechanism to uncouple co-produced miRNAs from each other. Overall, our findings indicate that the regulatory influence of TDMD in mammalian biology is widespread and posit the existence of many yet-unidentified transcripts that trigger miRNA degradation.

## Introduction

MicroRNAs (miRNAs) are ∼22-nucleotide (nt) RNAs that associate with Argonaute (AGO) proteins to guide the repression of mRNAs. Within the miRNA–AGO complex, the miRNA recognizes target RNAs through base-pairing interactions, primarily to the miRNA seed region (miRNA nucleotides 2–8), while the AGO protein recruits deadenylases that accelerate deadenylation of the targeted mRNA, which typically promotes mRNA decay (Jonas and Izaurralde, 2015; Bartel, 2018; Eisen et al., 2020). miRNA-directed regulation is both pervasive and biologically important, in that each of the 90 most broadly conserved miRNA families has, on average, >400 preferentially conserved targets (Friedman et al., 2009), and for most of these families, loss of function in mice results in physiological or developmental abnormalities that often severely impact fitness (Bartel, 2018).

In spite of the well-established regulatory logic governing canonical miRNA–target interactions, in certain cases, the direction of this logic is reversed. In these cases, when a target RNA engages both the seed of the miRNA and its 3′ region through extensive base pairing, the miRNA becomes destabilized (Ameres et al., 2010; Cazalla et al., 2010; Libri et al., 2012; Marcinowski et al., 2012; Xie et al., 2012; Lee et al., 2013; de la Mata et al., 2015; Bitetti et al., 2018; Ghini et al., 2018; Kleaveland et al., 2018; Sheu-Gruttadauria et al., 2019). A biological role for this phenomenon of target-direct miRNA degradation (TDMD) was first observed during infection of primate T-cells by a gamma-herpesvirus, which expresses a noncoding transcript, HSUR1, that directs degradation of host miR-27, a miRNA that might otherwise limit viral replication (Cazalla et al., 2010). Since then, TDMD has been found to be exploited by other herpesviruses that express unrelated transcripts that trigger degradation of specific host miRNAs (Libri et al., 2012, Marcinowski et al., 2012, Lee et al., 2013). More recently, cellular transcripts that direct degradation of endogenous miRNAs have also been discovered. In a founding example of this endogenous TDMD, a site within the mouse *Nrep* mRNA directs degradation of miR-29b, which shapes behavior of the animal, as does a site in an orthologous long noncoding RNA (lncRNA) in zebrafish (Bitetti et al., 2018). In another example, noted for its potency, a site within the Cyrano lncRNA directs degradation of miR-7, which reduces the level of miR-7 by >97% in some mouse tissues, including the cerebellum (Kleaveland et al., 2018). Two other examples of endogenous TDMD have been reported from studies using cultured mammalian cells (Ghini et al., 2018; Li et al., 2021), and another six have been identified in Drosophila cells or embryos (Kingston et al., 2022; Sheng et al., 2023). For instance, a site within the *Drosophila melanogaster* lncRNA Marge directs degradation of members of the miR-310 family, which is required for proper development of the embryonic cuticle (Kingston et al., 2022).

TDMD requires a Cullin-RING E3 ubiquitin ligase complex containing the substrate adapter ZSWIM8 (Han et al., 2020; Shi et al., 2020). In the current model of TDMD, this ZSWIM8 E3 ligase recognizes a distinct conformation assumed by the AGO–miRNA complex when extensively paired to a trigger site (Sheu-Gruttadauria et al., 2019), which results in the polyubiquitination and proteasomal destruction of AGO, exposing the miRNA to cellular nucleases (Han et al., 2020; Shi et al., 2020).

The loss of ZSWIM8 not only causes increased accumulation of known TDMD substrates, such as miR-7 and miR-29b in mammalian cells, but it also causes increased accumulation of other miRNAs, implicating these additional ZSWIM8-sensitve miRNAs as potential TDMD substrates. Thus far, 34 ZSWIM8-sensitve miRNAs have been identified in mouse or human cells (Shi et al., 2020; Li et al., 2022), 21 have been identified in Drosophila S2 cells or embryos (Shi et al., 2020; Kingston et al., 2022), and 10 have been identified in *C. elegans* gravid adults (Shi et al., 2020). These findings suggest that the scope of endogenous TDMD is broad, and that this pathway provides a conserved mechanism by which diverse animal species shape the levels of their miRNAs. Indeed, the influence of ZSWIM8 quantitatively explains the short half-lives of most short-lived miRNAs in both mouse embryonic fibroblasts (MEFs) and Drosophila S2 cells (Shi et al., 2020).

Interestingly, Dorado (Dora), the ZSWIM8 ortholog in flies, is essential for viability (Kingston et al., 2022), but EBAX-1, the ortholog in *C. elegans*, is not essential (Wang et al., 2013). Perhaps TDMD mediates divergent biological functions in different animals, presumably through variation in the cohort of miRNAs targeted for degradation. Another possibility is that other roles of ZSWIM8 contribute to these different phenotypic outcomes. Indeed, mechanisms unrelated to miRNAs are reported for EBAX-1/ZSWIM8-mediated promotion of proper neural development in *C. elegans* and mouse (Wang et al., 2013; Wang et al., 2022) and Dora-mediated promotion of proper hair formation in *D. melanogaster* (Molina-Pelayo et al., 2022). Here, we find that in mice, ZSWIM8 is required for lung sacculation and proper heart development, with loss of function causing perinatal lethality, accompanied by increased levels of more than 50 miRNAs from more than 40 different miRNA families.

## Results

### ZSWIM8 is required for perinatal viability

To assess the biological functions of ZSWIM8 in mice, we generated *Zswim8* loss-of-function mutant alleles using Cas9, targeting the second exon (Fig. 1A). This exon encodes the conserved BC-box and Cullin-box domains thought to be critical for ZSWIM8 activity as part of a Cullin-RING ligase (Wang et al., 2013), and targeting this exon abrogates TDMD in cell culture (Shi et al., 2020). Four independent mouse lines were recovered, each bearing a frameshifted allele predicted to produce truncated protein with disrupted function (Fig. 1A). As heterozygotes (*Zswim8*^+/–^), these mutant mice were grossly normal and fertile. We did not observe any phenotypic differences between the four lines bearing these alleles, and hence they were combined and treated as identical.

**Figure 1.**
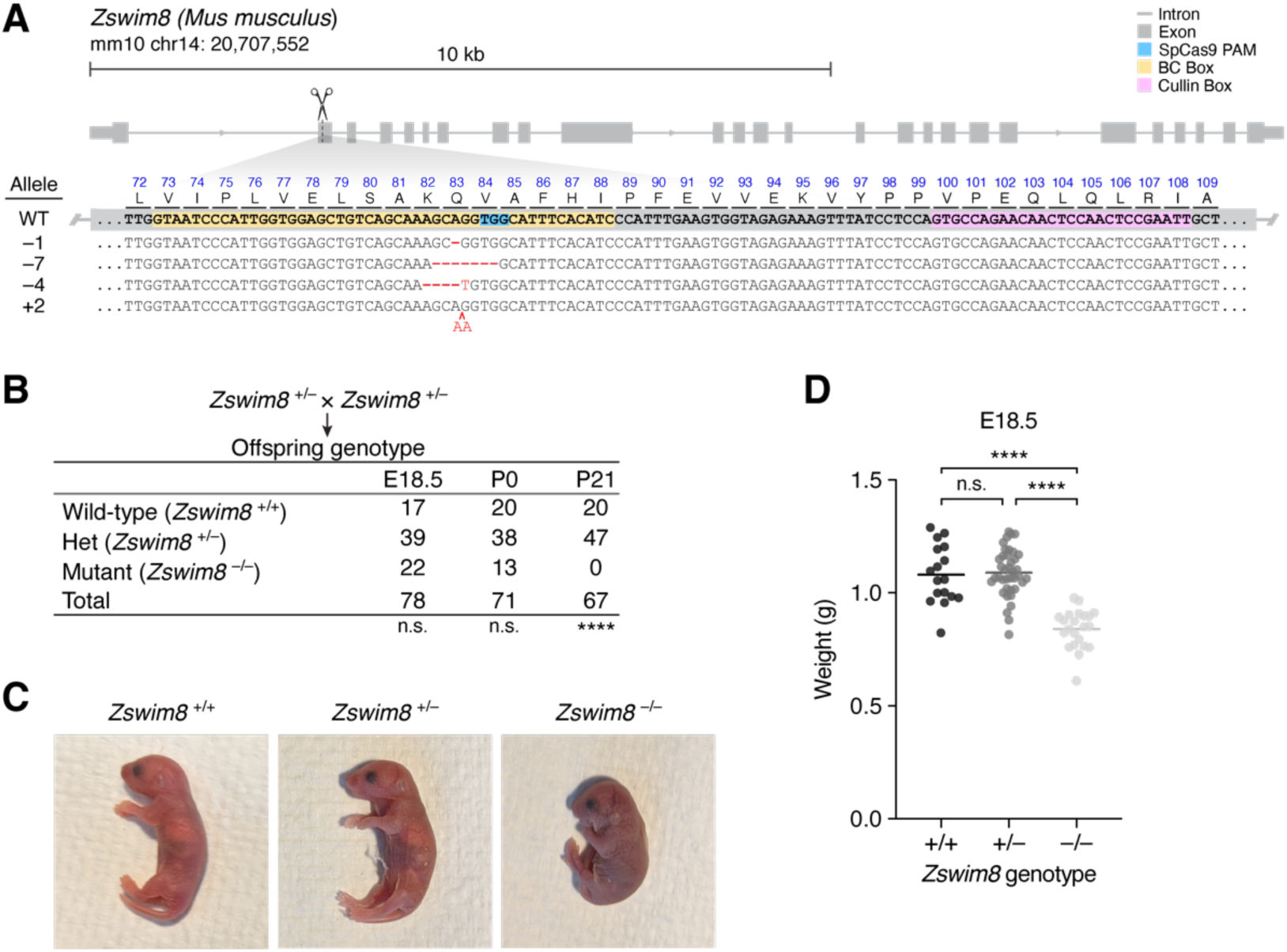
Gross phenotypes of *Zswim8^−/−^*mice. (A) Organization of the murine *Zswim8* genomic locus. Shown below the gene model (thick gray boxes, exonic coding sequence; thin gray boxes, exonic untranslated regions; gray lines, introns; blue box, Cas9 PAM; yellow box, annotated BC Box motif; pink box, annotated Cullin Box motif) are the genomic DNA sequences of the wild-type (WT) allele and four mutant alleles (“–1”, “–7”, “–4”, and “+2), as well as the amino acid sequence of the expected WT translation product. Blue text, WT protein sequence; black text, WT DNA sequence; red text, insertions; red dashes, deletions. (B) Penetrant lethality of *Zswim8*^−/−^ animals. Shown are genotypes of offspring produced from intercrosses between *Zswim8*^+/–^ parents. Offspring were counted at either embryonic E18.5, P0, or P21. Differences from the Mendelian expectation were evaluated using the Chi-square test (two-tailed; n.s., not significant, *p* > 0.05; ****, *p* < 0.0001). (C) Cyanotic phenotype of *Zswim8*^−/−^ pups. Shown are P0 neonate siblings with the indicated genotypes, cropped from the same photograph. (D) Smaller size of *Zswim8*^−/−^ embryos. Plotted are weights of E18.5 embryos produced from intercrosses between *Zswim8*^+/–^ parents. Horizontal lines indicate the mean. Significance of differences were evaluated using ANOVA (Tukey’s multiple comparisons test; n.s., not significant, *p* > 0.05; ****, *p* < 0.0001).

After intercrossing *Zswim8*^+/–^ mice and genotyping at the age of weaning (around postnatal day 21, P21), the ratio of heterozygous to wild-type offspring was within the Mendelian expectation, but no homozygous mutants (*Zswim8*^−/−^) were observed (Fig. 1B). In contrast, on the day of birth (P0) or at embryonic day 18.5 (E18.5), the expected Mendelian ratios of the three *Zswim8* genotypes were observed (Fig. 1B), indicating that *Zswim8*^−/−^ animals died either perinatally or during post-embryonic development.

Closer examination of *Zswim8*^−/−^ P0 neonates yielded several additional observations: 1) many, but not all, were dead at the time of observation; 2) nearly all, whether alive or dead, appeared cyanotic (Fig. 1C); 3) those that were alive at the time of observation died within several hours, often following a period of agonal breathing (Video S1), and 4) most appeared smaller than their littermates. In contrast, wild-type and *Zswim8^+/–^* neonates appeared normal and did not display any of the aforementioned phenotypes.

To quantify the size difference observed for *Zswim8*^−/−^ mice, we weighed embryos dissected at E18.5, approximately a day before their expected birth, to prevent potentially confounding differences in feeding. Whereas wild-type and *Zswim8*^+/–^ embryos did not significantly differ in weight, *Zswim8*^−/−^embryos were about 22% lighter (Fig. 1D).

### ZSWIM8 is required for proper embryonic development of heart and lung

The cyanotic and respiratory-distress phenotypes of *Zswim8*^−/−^ neonates suggested that these animals failed to achieve proper oxygenation after birth. Although such a phenotype could arise from defects in a number of different physiological processes, we decided to first examine the developmental anatomy of the cardiovascular and pulmonary systems.

Serial transverse sections of hearts at E18.5 revealed the presence of ventricular septal defects (VSDs) in three of four *Zswim8*^−/−^ embryos examined, but not their *Zswim8*^+/–^ littermates (Fig. 2A). VSDs are one of the most commonly recognized congenital heart defects and sometimes cause cyanosis in human patients (Mavroudis 2012). Although the VSDs might contribute to the cyanosis observed in *Zswim8^−/−^* newborns, they did not appear to be especially severe, raising the question of whether they were sufficient to explain the highly penetrant lethality. To search for additional defects that might contribute, we examined lung sections from E18.5 embryos that had never breathed air. Relative to lungs of *Zswim8*^+/–^ littermates, lungs of *Zswim8*^−/−^ embryos had a modest but significant reduction in total airspace (Fig. S1).

**Figure 2.**
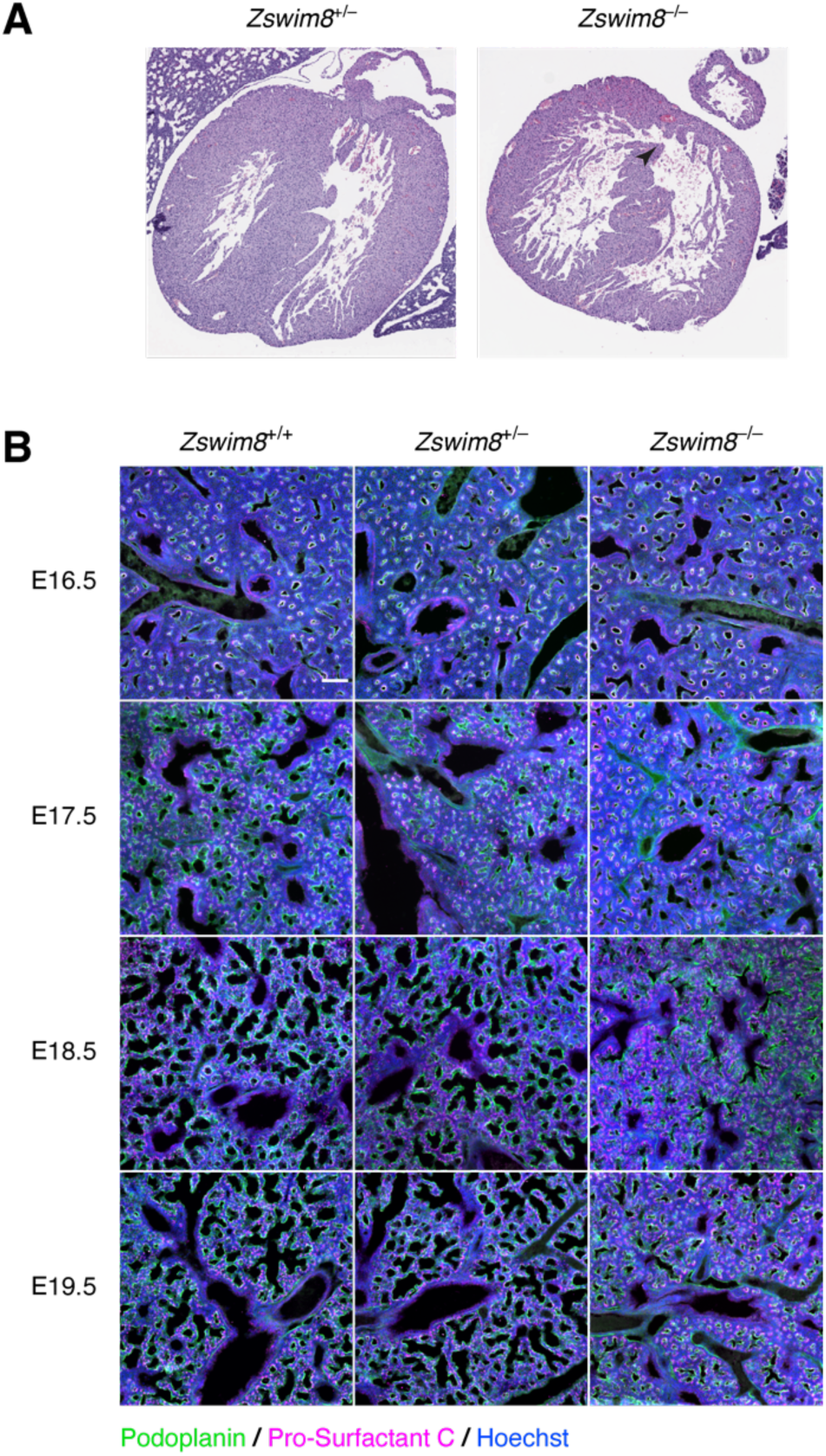
Tissue-level phenotypes of *Zswim8*^−/−^ embryos. (A) Heart defect in *Zswim8*^−/−^ embryos. Shown are representative images of H&E-stained transverse sections of hearts from *Zswim8*^+/–^ and *Zswim8*^−/−^ littermates at E18.5. Arrowhead points to a VSD. (B) Sacculation defect in *Zswim8*^−/−^ embryos. Shown are lung sections from *Zswim8*^+/+^, *Zswim8*^+/–^, and *Zswim8*^−/−^ embryos at the indicated developmental times immunostained for Podoplanin and Pro-surfactant C (green and magenta, respectively), and stained for DNA (Hoechst, blue). Scalebar represents 100 μm.

The embryonic lung develops initially as buds arising from the anterior foregut endoderm to form the trachea at ∼E9.5, which then extend into the surrounding mesenchyme and undergo a successive branching process to form the airway arbor (Herriges and Morrisey, 2014). During this branching morphogenesis, cell lineages develop along a proximal–distal axis, resulting in the formation of structural airways that terminate in distal tips. During the saccular phase of development, which begins at ∼E18.5, these distal tips undergo airspace expansion to form saccules that are precursors to alveoli, the future sites of gas exchange. These developing alveoli are populated by the two major alveolar epithelial cell lineages: the alveolar type I (AT1) cells, which are squamous and form most of the gas-exchange surface area, and alveolar type II (AT2) cells, which produce and secrete pulmonary surfactant (Morrisey and Hogan, 2010).

To examine the effect of ZSWIM8 loss on sacculation, we stained developing lungs from wild-type, *Zswim8*^+/–^, and *Zswim8*^−/−^ embryos, for Podoplanin and Pro-surfactant C—markers of AT1 and AT2 cells, respectively. As previously described for the saccular stage of development, we observed that wild-type lungs exhibited alveolar epithelial cell differentiation by E17.5 and the onset of airspace expansion at the distal tips at E18.5 (Wang et al., 2016; Morrisey and Hogan, 2010). Whereas lungs from wild-type and *Zswim8*^+/–^embryos appeared similar to each other throughout E16.5–E18.5, those from *Zswim8*^−/−^ embryos visibly diverged at E18.5, failing to undergo airspace expansion despite possessing approximately normal proportions of AT1 and AT2 cells (Fig. 2B). Notably, this airspace-expansion defect was localized to the alveolar compartment, as airways, which do not stain positive for Podoplanin, appeared grossly normal across all genotypes (Fig. 2B). The airspace defect in *Zswim8*^−/−^ lungs persisted to E19.5 (Fig. 2B), suggesting that proper saccular development was impeded through the time of birth.

### ZSWIM8 is required for proper maturation of alveolar epithelial cells

To more closely examine the cellular basis for the airspace-expansion defect, we dissociated whole lungs from two *Zswim8*^+/–^ and three *Zswim8*^−/−^ E18.5 embryos and performed single-cell RNA sequencing (scRNA-seq). After filtering for quality, data from 19,732 cells were captured and analyzed by UMAP embedding (McInnes et al., 2018) and unsupervised clustering.

For clusters representing most of these lineages, the two genotypes were similarly distributed (Fig. 3A). A notable exception was the cluster corresponding to the lung epithelial lineage—characterized by expression of *Nkx2.1* (Herriges and Morissey, 2014) (Fig. S2A), in which cells from the *Zswim8*^−/−^ samples were depleted from the periphery of the embedded space (Fig. 3B). Re-embedding and re-clustering of this compartment revealed regions with high expression of *Ager* and *Sftpb,* canonical markers for AT1 and AT2 cells, respectively (Fig. 3B), with peripheral sub-regions possessing near-exclusive expression of one or the other. These sub-regions likely represented populations of the more developmentally mature forms of these cell types, as evident from enriched expression of other canonical marker genes within the corresponding clusters (Clusters 3, 5 and 0; Fig. S2D). Cells from *Zswim8*^−/−^ lungs were depleted from these sub-regions, concentrating instead in intervening clusters (clusters 1, 7, and 8), in which they substantially outnumbered cells of *Zswim8*^+/–^ lungs (Fig. 3B, S2B). Of these, clusters 1 and 7 were the more populated, and had intermediate expression of both canonical AT1 and AT2 lineage markers. Notably, cluster 1 appeared to form a bridge in the embedded space between the AT1 and AT2 regions, and two of its top four most significant marker genes, *Ccn1* and *H19* (Fig. S2D), were among the top five marker genes for an AT1 precursor state normally present at E17.5 (Frank et al., 2019) (Fig. 3C). Cluster 7 was uniquely characterized by a high proportion of cells in the G2/M phase (Fig. S2C) and significantly enriched expression (*p* < 10^−5^) of four out of five AT1 precursor markers, as well as for four out of five AT2 precursor markers (Frank et al., 2019) (Fig. 3C). The expression patterns of these precursor markers generally corresponded well with our clustering (Fig. S2E).

**Figure 3.**
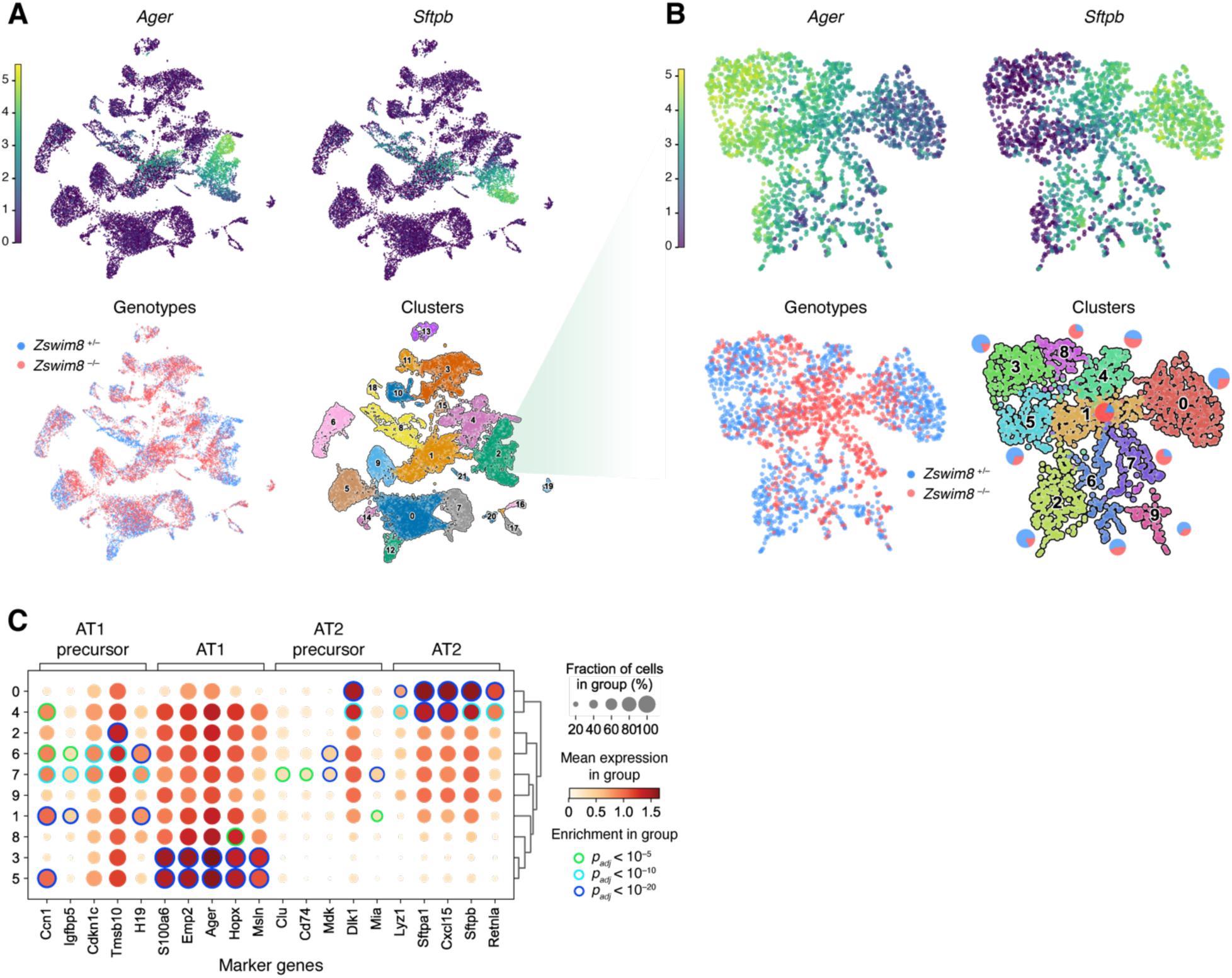
Improper maturation of epithelial cells from lungs of *Zswim8*^−/−^ embryos. (A) UMAP analysis of scRNA-seq data from all cells recovered from whole lungs of E18.5 embryos. Each point shown represents a cell from either *Zswim8^+/–^*(*n* = 2), or *Zswim8^−/−^* (*n* = 3) embryos. Top: coloring indicates expression of indicated genes; color bar is in units of ln(1+ CP10K). CP10K, counts per ten thousand unique counts. Bottom-left: Coloring indicates genotype, with cells from embryos of the same genotype pooled together (blue, *Zswim8^+/–^*; red, *Zswim8^−/−^*). Bottom-right: Coloring indicates cluster, with cluster numbers labeled and further characterized in Fig. S2A. (B) Re-embedded UMAP plots showing cells from Cluster 2 of panel (A). Pie charts in the bottom-right indicate percentages of cells of each genotype in each cluster. Otherwise, this panel is as in (A). (C) Expression of marker genes (column labels) reported for AT1 precursor, AT1, AT2 precursor, and AT2 cells (Frank et al., 2019) in clusters identified in (B). Row labels correspond to cluster numbers from (B). Size of discs indicates the fraction of cells in the cluster with detectable counts for the gene. Fill color of discs indicates mean expression in the cluster; color bar is in units of ln(1+ CP10K). Edge color of discs indicates statistical significance of enrichment for the gene in that cluster, relative to all clusters shown, as evaluated by the Wilcoxon rank-sum test, adjusted by the Benjamini-Hochberg method (green, *p_adj_* < 10^−5^; light blue, *p_adj_* < 10^−10^; dark blue, *p_adj_* < 10^−20^).

Together, these results supported a model in which the sacculation defect observed in *Zswim8*^−/−^ lungs resulted from a failure of AT1 and AT2 precursors to properly differentiate into their respective mature forms after E17.5, causing the aberrant persistence and/or proliferation of these precursors at E18.5, and a block in sacculation.

### ZSWIM8 has a widespread impact on embryonic miRNA levels

To investigate the impact of ZSWIM8 on miRNA levels, we performed small-RNA sequencing (sRNA-seq) on samples from forebrain, hindbrain, eye, heart, lung, liver, stomach, kidney, intestine, skin, skeletal muscle, and placenta dissected from *Zswim8*^+/–^ and *Zswim8*^−/−^ E18.5 embryos. This late embryonic stage was chosen to capture developmental differences in gene expression between genotypes, while avoiding secondary effects arising from the need of newborns to breath air.

The abundance of miRNAs quantified for each tissue of the same genotype correlated well across two replicates (Fig. S3A). To identify miRNAs that were significantly up-regulated in *Zswim8*^−/−^ tissues, we used a statistical approach developed for detecting biological changes that are expected to be unidirectional, based on a bi-beta-uniform mixture (BBUM) model, which offers increased robustness to secondary effects, as well as false-discovery rate (FDR)-adjusted significance thresholds (Wang et al., 2022). Across the twelve tissues examined, this approach generally improved sensitivity in identifying miRNAs that increased upon ZSWIM8 loss (Fig. S3B).

The miRNAs that were designated as significantly upregulated by BBUM analysis (FDR-adjusted *p* value < 0.05) were further analyzed based on the behavior of each miRNA relative to that of its passenger strand—i.e., the strand concurrently processed from the other arm of the miRNA precursor during the normal course of miRNA biogenesis (Bartel, 2018). Because TDMD occurs after the miRNA associates with AGO and dissociates from its passenger strand, authentic TDMD substrates increase upon loss of ZSWIM8, without a corresponding increase in their passenger strands (de la Mata et al., 2015). Accordingly, the ZSWIM8-sensitive miRNAs of each tissue were filtered to remove those that did not increase significantly more than their passenger strands upon *Zswim8* knockout. These miRNAs that were removed from the main set of ZSWIM8-sensitive miRNAs were designated as indirectly ZSWIM8-sensitive miRNAs. Although we cannot rule out the possibility that for some of these miRNAs both the miRNAs and their passenger strands were ZSWIM8 sensitive, we excluded them from the main set of ZSWIM8-sensitive miRNAs with the idea that their increased levels observed upon *Zswim8* knockout were most likely the result of increased pri-miRNA transcription or processing, or some other secondary effect of ZSWIM8 loss that increased both the miRNAs and their passenger strands.

After applying these criteria to identify the ZSWIM8-sensitive miRNAs in each of the 12 tissues, we compared the results for each miRNA in different tissues to search for evidence of false-positives and false-negatives. When comparing across tissues, a few miRNAs appeared to have passed the annotation thresholds in a few individual tissues (typically one) due to either transcription effects or extreme variability; these were removed from our set of ZSWIM8-sensitive miRNAs as suspected false-positives. Another nine miRNAs did not meet our requirement for annotation as ZSWIM8-sensitive in any single tissue, yet were nonetheless broadly upregulated (median log_2_ fold change in *Zswim8*^−/−^ >0.2), with increased levels exceeding those of their respective passenger strands broadly (in at least 11 out of 12 tissues) and substantially (median log_2_ magnitude >0.2); these nine broadly sensitive miRNAs were considered false negatives of our analyses of individual tissues, and were added to our list of ZSWIM8-sensitive miRNAs of E18.5 embryos, bringing the total to 51 different miRNAs from 43 different miRNA families (Table S1).

Among the 42 miRNAs classified as ZSWIM8-sensitive in one tissue, most were independently classified as ZSWIM8-sensitive in other tissues (Fig. S3C). In many other cases, miRNAs classified as ZSWIM8-sensitive in at least one tissue increased upon ZSWIM8 loss in additional tissues, but not to the extent required to passed the significance threshold for independent classification in those tissues. For those cases in which this elevation significantly exceeded that observed for the passenger strand, we anticipate that with more sensitive analyses (e.g., with more biological replicates), the miRNAs will ultimately be confidently classified as ZSWIM8-sensitive miRNAs in the broader range of tissues. In the meantime, we designated them as marginally ZSWIM8-sensitive in those tissues (Fig. 4A, S3D; Table S1).

**Figure 4.**
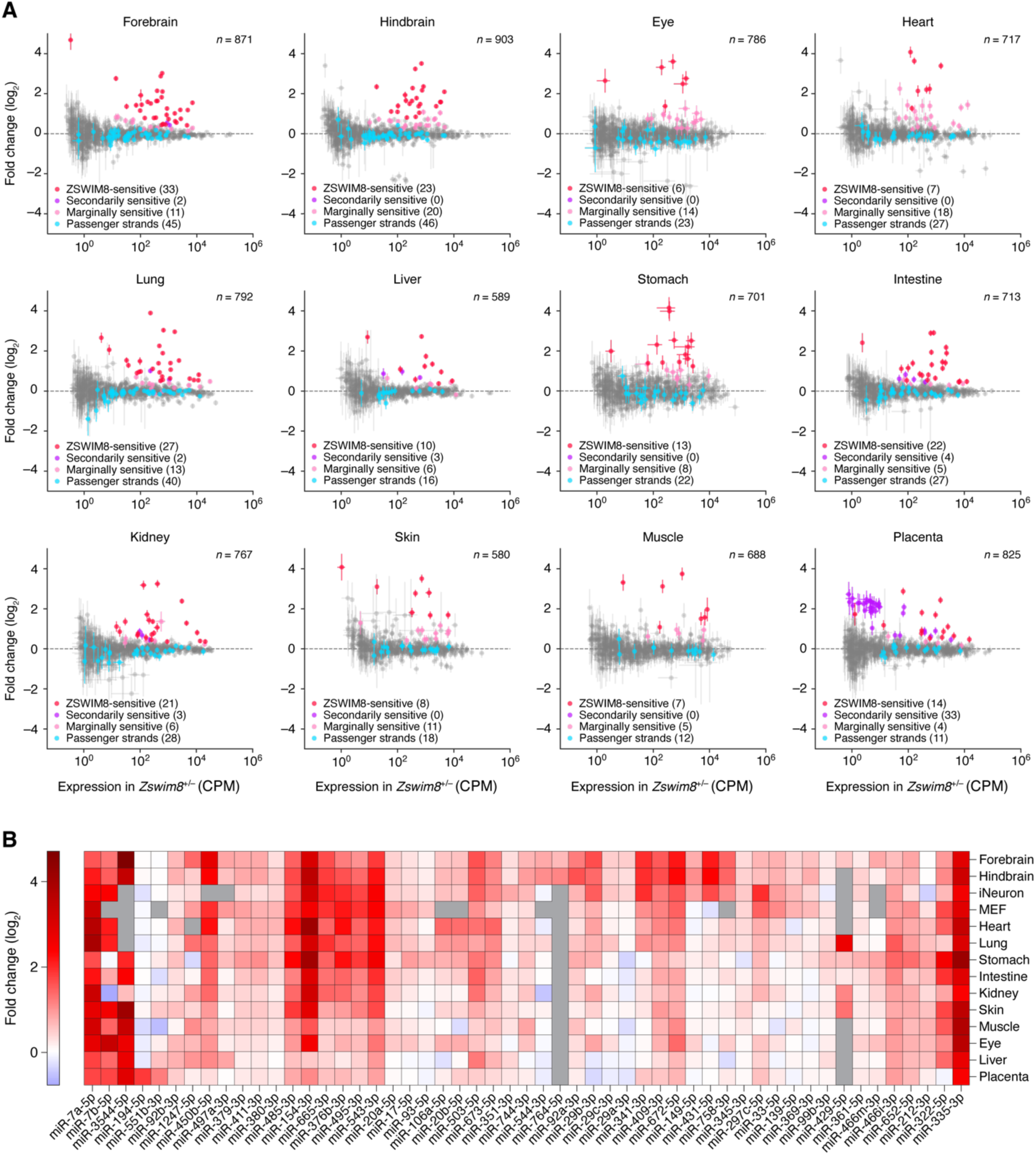
The impact of ZSWIM8 on miRNA levels in embryonic tissues. (A) The influence of ZSWIM8 on miRNA levels in the indicated tissues of mouse embryos, as determined by sRNA-seq. Plotted are fold changes in miRNA (or passenger strand) levels observed when comparing the results for tissues from *Zswim8*^−/−^ E18.5 embryos with those from *Zswim8*^+/–^ E18.5 embryos (error bars, standard error of two biological replicates). Red denotes ZSWIM8-sensitive miRNAs, purple denotes indirectly ZSWIM8-sensitive miRNAs (and their passenger strands), pink denotes marginally ZSWIM8-sensitive miRNAs, blue denotes passenger strands of sensitive and marginally sensitive miRNAs, and gray denotes all other annotated miRNAs or passenger strands that exceeded our expression threshold. CPM, counts per million. *n*, total number of small RNAs analyzed. (B) The influence of ZSWIM8 on miRNAs classified as ZSWIM8-sensitive in at least one of the twelve embryonic tissues. The heatmap indicates fold-changes (key) observed when comparing the results for tissues from *Zswim8*^−/−^ E18.5 embryos with those from *Zswim8*^+/–^ E18.5 embryos (Table S1). Also included are results for these same miRNAs observed after polyclonal *Zswim8* knockout in MEFs and iNeurons (Shi et al., 2020), as well as results for two miRNAs that were called as ZSWIM8-sensitive in at least one of those cell lines and were marginally ZSWIM8-sensitive in at least one embryonic tissue (miR-93-5p and miR-297c-5p). Gray squares indicate contexts in which the number of miRNA reads did not exceed the detection threshold of 5 CPM in each library prepared from the corresponding tissue.

### ZSWIM8 can change the dominant miRNA strand or isoform

For nine miRNA duplexes (miR-335, miR-429, miR-466i, miR-497a, miR-544, miR-652, miR-764, miR-744, and miR-99b), the strand annotated as the passenger strand was ZSWIM8-sensitive (Table S1). In these cases, we considered the ZSWIM8-sensitive strand as the miRNA and the other strand as the passenger strand for purposes of evaluating ZSWIM8 sensitivity. The miR-744 duplex was noteworthy in this respect, as its annotated passenger strand (miR-744-3p) was called as ZSWIM8-sensitive based on its broad albeit modest sensitivity in 11 of 12 tissues, whereas its annotated miRNA strand (miR-744-5p) is called as ZSWIM8-sensitive in induced mouse neurons (iNeurons) (Shi et al., 2020).

For five miRNA duplexes (miR-154, miR-335, miR-411, miR-450b, and miR-532), the strand that is normally less abundant became the more abundant strand upon loss of ZSWIM8 in a least one of the 12 tissues (Fig. 5A). For each of these miRNAs except miR-335, both strands were annotated as guides. Thus, tissue-selective ZSWIM8 sensitivity could explain the previous report of miR-154 “arm switching” (Chiang et al., 2010), a phenomenon in which the more abundant strand of a duplex differs in some contexts compared to others. Likewise, for miR-450b, the greater ZSWIM8 sensitivity of the 5p isoform in some tissues (e.g., eye) than in others (e.g., placenta) led to a newly identified example of arm switching (Fig. 5A).

**Figure 5.**
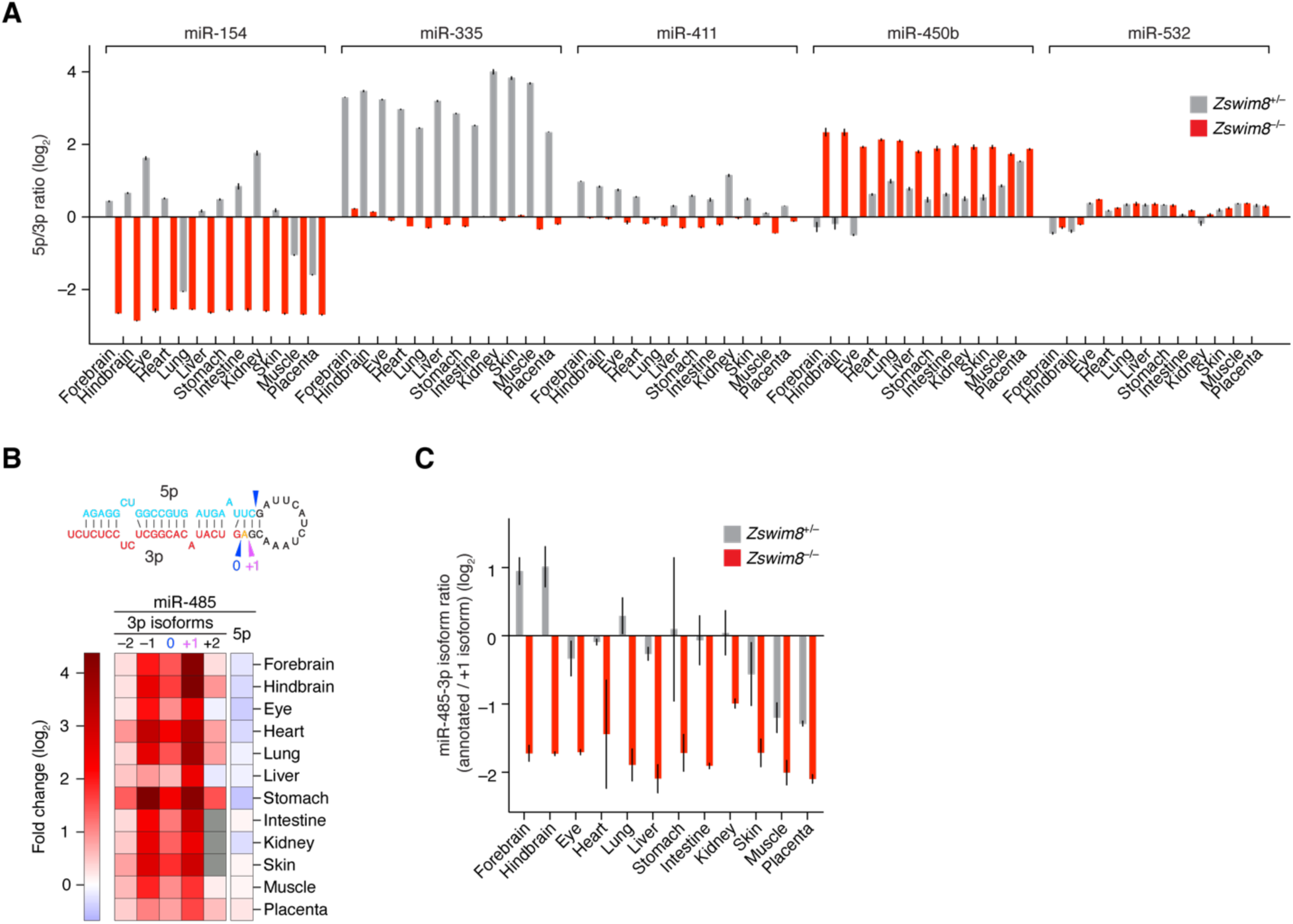
Influence of ZSWIM8 on miRNA isoform abundance and arm switching. (A) The influence of ZSWIM8 on the relative abundances of the two strands of miRNA duplexes. Plotted are ratios of the mean levels of 5p and 3p strands for the indicated miRNA duplexes in the indicated tissues, as quantified by sRNA-seq (error bars, error propagated from standard error of two biological replicates). (B) The influence of ZSWIM8 on the abundances of miR-485-3p isoforms in the indicated tissues of mouse embryos, as determined by sRNA-seq. The heatmap indicates fold-changes (key) observed when comparing the results for tissues from *Zswim8*^−/−^ E18.5 embryos with those from *Zswim8*^+/–^ E18.5 embryos, as in Fig. 4B. The –2, –1, 0, +1, and +2 labels indicate 5′ isoforms of miR-485-3p, with –1 and –2 denoting isoforms with 1- and 2-nt 5′ truncations, respectively, +1 denoting an isoform with 1-nt templated extension (miR-485-3p.2), and +2 denoting an isoform with 2-nt templated extension—all relative to the annotated isoform (miR-485-3.1p), denoted as 0. The inferred Dicer-processing sites that generated the annotated (0) and +1 miR-485-3p isoforms (as well as the annotated miR-485-5p strand) from the miR-485 precursor are indicated with arrowheads in the schematic above the heatmap. The annotated and +1 isoforms are emphasized because these were the two 3p isoforms with highest abundance across all tissues examined. (C) The influence of ZSWIM8 on the relative abundances of the annotated and +1 isoforms of miR-485-3p. Plotted are ratios of the two isoforms in *Zswim8*^+/–^ and *Zswim8*^−/−^ embryonic tissues, as quantified by sRNA-seq (error bars, error propagated from standard error of two biological replicates).

In a related result, an isoform of miR-485-3p, which had an additional nucleotide at its 5′ end (a “+1” isoform, which we call miR-485.2-3p) and was previously unannotated by TargetScan7 (Agarwal et al., 2015), was even more ZSWIM8-sensitive than the annotated isoform (miR-485.1-3p) (Fig. 5B). As a result, upon ZSWIM8 loss, miR-485.2-3p accumulated to levels higher than those of either annotated guide strands (miR-485-5p and miR-485-3p) in each tissue examined (Fig. 5C).

Taking the union of the 51 unique miRNAs identified as ZSWIM8-sensitive in our analyses of the twelve E18.5 tissues with the 29 miRNAs previously identified as ZSWIM8-sensitive in cultured MEFs and iNeurons (Shi et al., 2020) brought the total number of ZSWIM8-sensitive miRNAs identified in mouse cells to 58, representing 48 miRNA families. A summary of the ZSWIM8-sensitivity profiles for the 53 miRNAs found to be at least marginally sensitive in embryonic tissues revealed distinct, context-specific patterns of sensitivity (Fig. 4B). For example, some miRNAs, such as miR-431-5p, were preferentially ZSWIM8-sensitive in brain or iNeuron samples despite being widely expressed, whereas others, such as miR-335-3p, were broadly and highly ZSWIM8-sensitive in every context examined. This pattern of sensitivity was unlikely to be explained by tissue-level differences in *Zswim8* mRNA expression, in that *Zswim8* mRNA levels were largely uniform across these tissues, although somewhat higher in brain and markedly lower, but still present, in liver (Fig. S4A). Interestingly, with the exception of placenta, *Zswim8* mRNA was slightly elevated in *Zswim8*^−/−^ tissues, hinting at feedback mechanisms governing its expression (Fig. S4A). In sum, when considering the marginally sensitive miRNAs together with the sensitive miRNAs, each tissue analyzed had at least 12 miRNAs that had ZSWIM8 sensitivity resembling that of TDMD substrates, with the numbers for brain and lung tissues exceeding 40. These results implied that TDMD has a widespread influence in shaping miRNA levels throughout the embryo.

### ZSWIM8-sensitive miRNAs are disproportionately produced from genomic clusters

Many miRNAs are produced from genomic clusters, wherein multiple hairpin-encoding miRNA genes are juxtaposed and often transcribed together in polycistrons. Such an arrangement enables co-expression of clustered miRNAs (Baskerville and Bartel, 2005) and can facilitate the processing of suboptimal miRNA hairpins (Fang and Bartel, 2020; Hutter et al., 2020; Shang et al., 2020). However, situations might arise in which decoupling the expression of cluster members could be advantageous, and TDMD might, in some cases, help enact this decoupling. As an initial exploration of this possibility, we examined the genomic distribution of the 58 miRNAs called as ZSWIM8-sensitive in mouse embryos or mouse cell lines, which corresponded to 61 miRNA hairpin loci. Of these 61 loci, 44 (or 72%) mapped to miRNA clusters (Fig. 6A), defined here as regions in which adjacent miRNA genes are separated by no more than 10 kilobases (kb). When considering that ∼46% of miRNAs derived from clusters, the fraction of ZSWIM8-sensitive miRNAs that derived from clusters was significantly higher than expected by chance (Fig. 6B).

**Figure 6.**
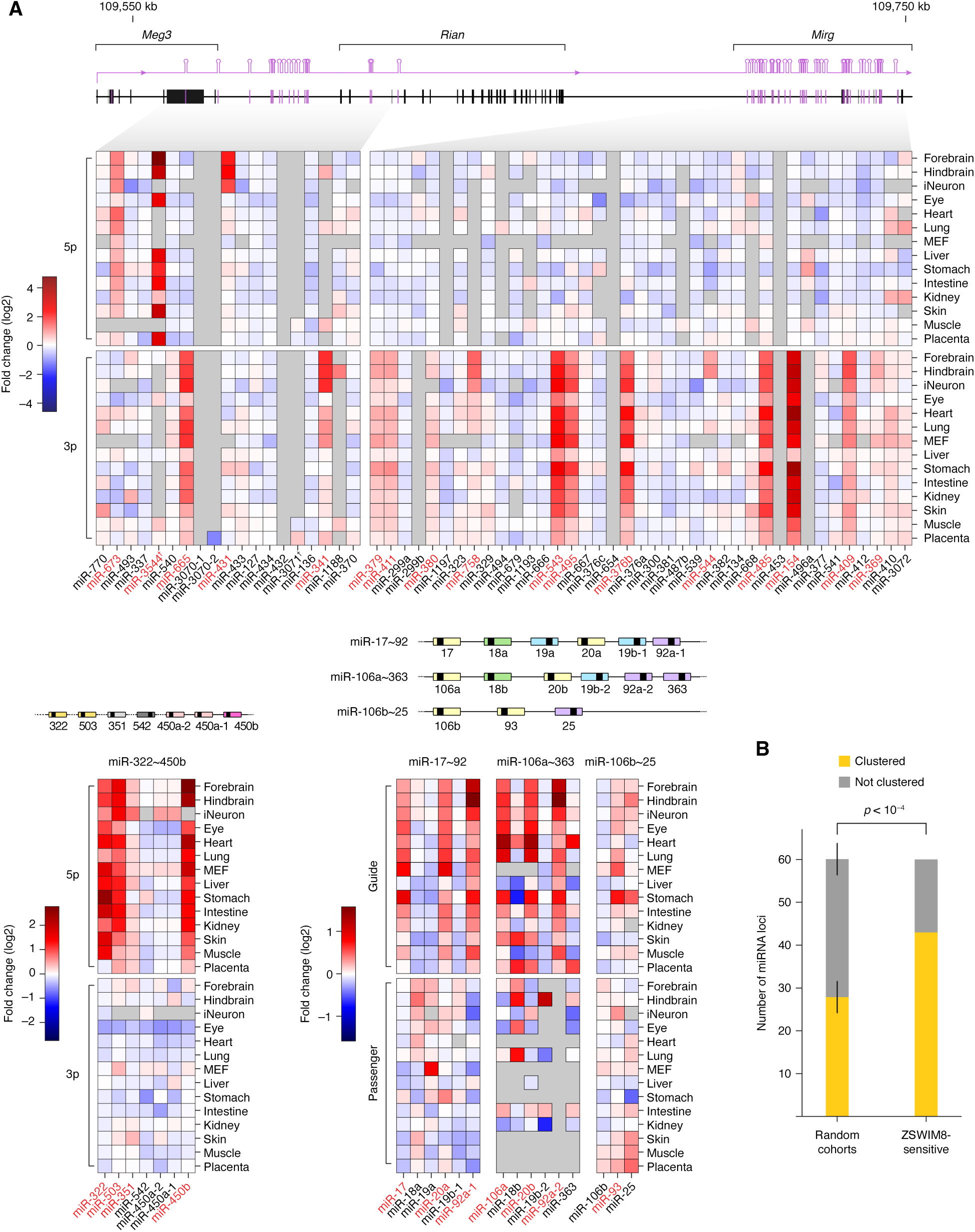
Genomic organization of ZSWIM8-sensitive miRNAs. (A) miRNA gene clusters producing a preponderance of ZSWIM8-sensitive miRNAs. Shown are fold changes in miRNA levels observed in *Zswim8*^−/−^ tissues, relative to *Zswim8*^+/–^ tissues from E18.5 embryos, as well as in MEFs and iNeurons (Shi et al., 2020). The miRNAs are ordered by position within the cluster, and organized by strand (either guide vs. passenger strand, as in the case of the miR-17∼92 and paralogous clusters, or 5p vs. 3p strand for the other clusters). Each value is the average of two biological replicates. Red text denotes ZSWIM8-sensitive miRNAs; dagger denotes miRNA apparently derived from the antisense strand of the paternal allele. (B) A tendency of ZSWIM8-sensitive miRNAs to derive from clustered miRNA genes. The bar plots show the proportion of miRNA loci produced from a miRNA cluster for the set of 61 loci encoding ZSWIM8-sensitive miRNAs, compared to random cohorts of the same size drawn without replacement from the set of all miRNA loci. Error bars show standard deviation across 10,000 random cohorts. The *p* value was calculated by hypergeometric test.

Strikingly, 17 ZSWIM8-sensitive miRNAs derived from the mammal-specific, imprinted *Dlk1-Dio3* locus on Chromosome 12, which encodes ∼116 mature miRNA strands produced from ∼58 hairpin loci, most of which are expressed from the maternal allele (Rocha et al., 2008) (Fig. 6A). Of these 17, 12 derived from the maternally expressed *Mirg* gene, which encodes 76 mature miRNA strands produced from 38 hairpin loci thought to be transcribed as a single poly-cistronic transcript. The remaining five derived from an upstream region of the *Dlk1*-*Dio3* locus, in the vicinity of the maternally expressed *Meg3* gene, although one of these, miR-3544-5p, which had the greatest sensitivity of all ZSWIM8-sensitive miRNAs (Fig. 4B, Table S1), derived from the opposite strand, several kb downstream of *Rtl1*, an imprinted gene that is predominantly expressed from the paternal allele (Seitz et al., 2003). Because many miRNAs of the *Dlk1-Dio3* locus are related to each other, multiple ZSWIM8-senstitive members might share a common trigger RNA. Supporting this idea, some pairs of miRNAs from this locus had similar patterns of sensitivity to ZSWIM8 across different tissue/cellular contexts (e.g., miR-495-3p and miR-543-3p; miR-379-3p and miR-411-3p) (Fig. 4B).

Other ZSWIM8-sensitive miRNAs derived from three paralogous clusters—the miR-17∼92 cluster, the miR-106a∼363 cluster, and the miR-106b∼25 cluster (Fig. 6A). Across the three paralogous clusters, most members of the miR-17 family (miR-17, miR-20a, miR-106a, miR-20b, and miR-93) were ZSWIM8-sensitive, the exception being miR-106b. Also sensitive was miR-92a, which is produced from both the miR-17∼92 and miR-106a∼363 clusters, although among the other four members of the miR-92 family (miR-92b, miR-25, miR-363, and miR-367), only miR-92b was also ZSWIM8 sensitive (Fig. 6A).

Other ZSWIM8-sensitive miRNAs included three miRNAs from the miR-322∼450b cluster (miR-322-5p, miR-503-5p, and miR-450b-5p). These were ZSWIM8-sensitive in nearly every context examined (Fig. 6A). Nonetheless, they each had different patterns of sensitivity, suggesting that different triggers acted upon them, even though miR-322-5p and miR-503-5p have similar seed sequences.

### ZSWIM8 reduces repression of targets of ZSWIM8-sensitive miRNAs

We next examined the consequences of these widespread changes in miRNA levels on the transcriptomes of tissues collected from *Zswim8^−/−^*embryos. For about half of the tissues, the effects were modest overall, with only a small number of individual mRNAs passing our significance threshold for differential expression (Fig. S4D). In heart, lung, liver, skin, and placenta, the number of differentially expressed mRNAs was greater, ranging in the tens to hundreds. The most widespread changes occurred in lung, presumably, in part, a consequence of perturbed lung epithelial development (Fig. 3B). In most tissues, more mRNAs significantly decreased than increased (Fig. S4D), suggesting that the overall increase in miRNA dosage driven by aberrant persistence of ZSWIM8-sensitive miRNAs could be causing some of these changes.

To assess this possibility, we examined predicted targets of the families of ZSWIM8-sensitive miRNAs whose members collectively increased the most upon ZSWIM8 loss (Fig. S4E). Across the twelve tissues, the most affected families varied in identity, with the most affected family increasing the total miRNA pool by 0.5–2% (Fig. S4E). For miR-7, the family most affected in forebrain, the levels of mRNAs predicted to be most susceptible to miR-7– mediated repression (i.e., the top predicted targets of miR-7) tended to decrease, albeit modestly, in the forebrain upon loss of ZSWIM8, as expected if the elevated levels of miR-7 caused increased repression of its regulatory targets (Fig. 7A; p < 0.001, comparing fold-changes of top predicted targets to those of control mRNAs with no canonical sites to miR-7, Wilcoxon’s rank-sum test). Evidence for enhanced repression was also detected among the larger set of mRNAs with conserved sites to miR-7 (conserved predicted targets, p < 0.05) and among the even larger set of mRNAs with any 7–8-nt canonical site to miR-7 (all predicted targets, p < 0.001)—albeit with even smaller median fold changes than that observed for the top predicted targets (Fig. 7A). These results resembled those observed upon mutation of the Cyrano lncRNA, which directs ZSWIM8-dependent degradation of miR-7 (Kleaveland et al., 2018; Han et al., 2020; Shi et al., 2020). Similar results were observed for the predicted targets of the most affect miRNA family in lung (miR-15) and in heart (miR-503) (Fig. 7A). Indeed, for 10 of the 12 tissues examined, analogous evidence for enhanced repression by the most affected miRNA was observed, and for most tissues, this evidence extended to the predicted targets of the second- and third-most affected miRNA families (Fig. 7B). These results indicated that loss of ZSWIM8 causes modest but widespread dysregulation of gene expression across many different organ systems, and suggested that these changes are driven, in part, by an overaccumulation of miRNAs presumed to be endogenous substrates of TDMD.

**Figure 7.**
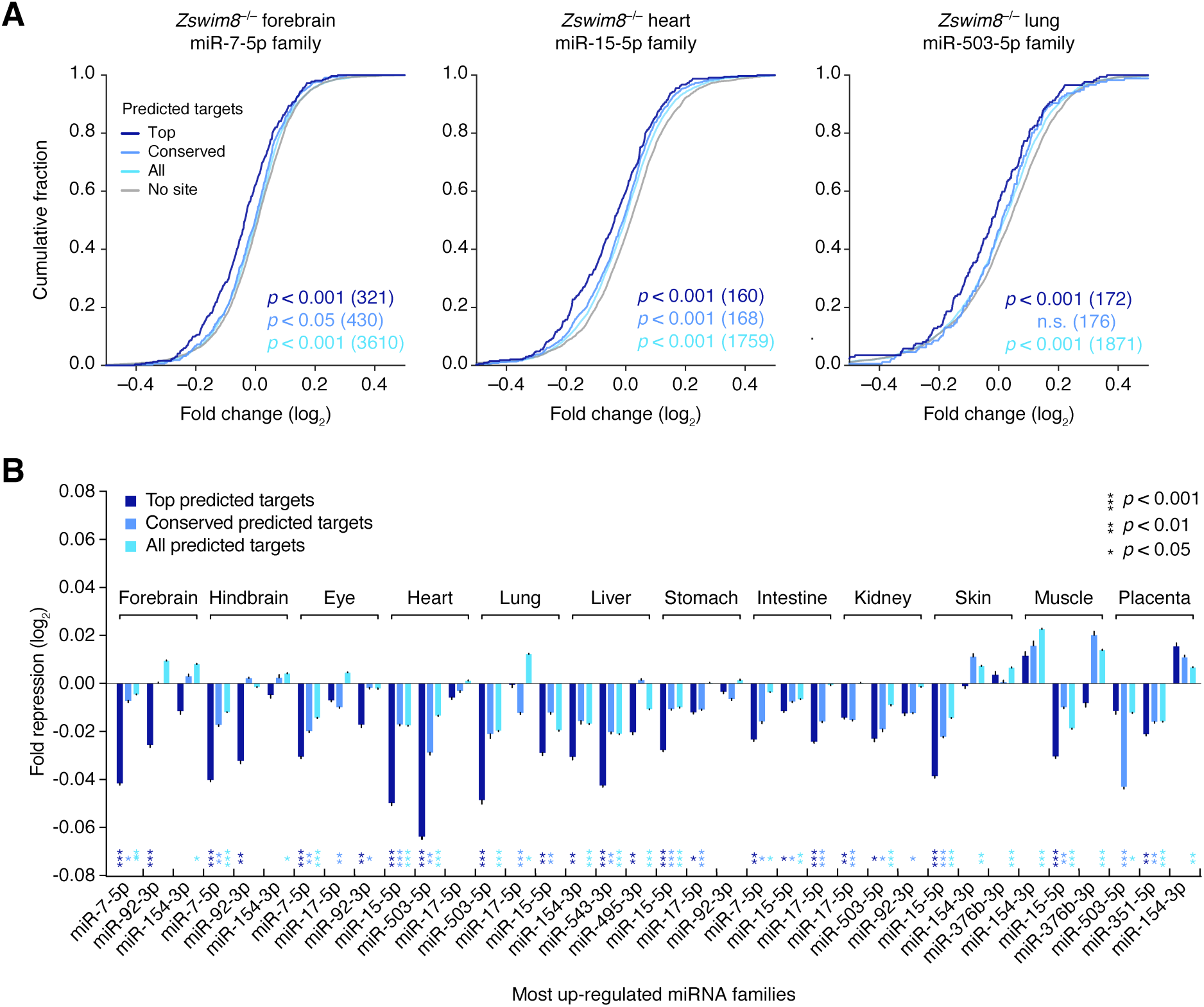
Increased repression of predicted targets of ZSWIM8-sensitive miRNAs upon loss of ZSWIM8. (A) Increased repression of predicted targets of the miRNA family whose members increase the most in embryonic forebrain (left), heart (middle), and lung (right). Plotted are cumulative distributions of mRNA fold changes in indicated tissues of *Zswim8*^−/−^ E18.5 embryos, relative to *Zswim8*^+/–^ embryos, for each of four sets of mRNAs: top predicted targets (top; dark blue), conserved predicted targets (conserved; medium blue), and all predicted targets (all; light blue) of the indicated miRNA family, as well as transcripts containing no canonical site to that family, which were length-matched to transcripts in the set of all predicted targets and randomly sampled at a five-to-one ratio (no site; gray). *p* values were calculated (Wilcoxon’s rank-sum test) comparing results for each set of predicted targets and its length-matched no-site cohort; for clarity, only the no-site cohort for the set of all predicted targets is shown. (B) Increased repression of predicted targets of the three miRNA families whose members collectively increased most in the indicated tissues from *Zswim8*^−/−^ embryos, relative to that of *Zswim8*^+/–^ embryos (excluding the eutherian miR-335-3p family, which evolved too recently to possess confidently predicted conserved targets (Friedman et al., 2009)). To summarize the degree of miRNA-mediated repression, the median fold change of the set of predicted targets was normalized to that of its length-matched no-site cohorts (error bars, standard error of the mean from 21 independent no-site cohorts). For each such sampling, a *p*-value was calculated as in (A), and the median *p*-value from all samples is shown for each family and each tissue, with coloring corresponding to the target set.

As expected, one of the transcripts reduced most upon loss of ZSWIM8 was the brain-enriched circular RNA *Cdr1as* (Fig. S4B). This circular RNA has 130 sites to miR-7 (Hansen et al., 2013; Memczak et al., 2013) and is the transcript most affected by mutation of Cyrano (Kleaveland et al., 2018). Interestingly, Cyrano was also sensitive to loss of ZSWIM8, decreasing in all 12 tissues examined from *Zswim8^−/−^*embryos (Fig. S4C), despite the fact that mutations that either reduce miR-7 or disrupt the site that normally directs ZSWIM8-dependent miR-7 degradation do not impact Cyrano levels in any tissues examined (Kleaveland et al., 2018). One potential explanation for these observations is that repression of Cyrano mediated by the intact site might normally be kinetically disfavored in comparison to TDMD, but in the absence of ZSWIM8, this repression has time to occur, as proposed for analogous cases in Drosophila in which the consequences of disrupting trigger sites have been compared to those of disrupting TDMD (Kingston et al., 2022). An alternative possibility is that other miRNAs accumulating in the absence of ZSWIM8 might mediate increased destruction of Cyrano. Indeed, within 100 bp of its miR-7 trigger site, Cyrano harbors an apparently functional regulatory site to the miR-92 family (Li et al., 2021), which includes miR-92a and miR-92b, two ZSWIM8-sensitive miRNAs that were broadly elevated across *Zswim8*^−/−^ tissues, especially the brain (Fig. 4B, S4E).

## Discussion

### Phenotypic consequences of ZSWIM8 knockout

Our finding that ZSWIM8 is essential for perinatal viability in mouse underscores the biological importance of this broadly conserved gene. The severity and penetrance of the *Zswim8* loss-of-function phenotype resembles that observed in *D. melanogaster*, where point mutants in the *Zswim8* ortholog *Dora* are lethal (90–100% penetrance, depending on the allele and genetic background), with most individuals dying during embryonic or early larval development (Kingston et al., 2022). In contrast, null mutants of the *Caenorhabditis elegans* ortholog*, ebax-1*, are viable, albeit with defects in axon guidance, locomotion, egg laying, and male mating (Wang et al., 2013), suggesting that ZSWIM8 plays varied biological roles across metazoa. The extent to which phenotypes observed in these different animals arise from the loss of TDMD, rather than some other function of ZSWIM8, is still an open question. Indeed, ZSWIM8 is proposed to regulate neural development, myogenesis, and actin dynamics through mechanisms that do not involve TDMD (Wang et al., 2013, Wang et al., 2022, Okumura et al., 2021, Molina-Pelayo et al., 2022).

In mice, the lung sacculation defect that we observed in *Zswim8*^−/−^embryos presumably caused the perinatal lethality, with perhaps some contribution from the heart VSD. Interestingly, at E18.5, the lungs of *Zswim8*^−/−^ embryos resembled those of *Hdac3* mutants, in which AT1 cells appear to have an autonomous spreading defect caused by elevated expression of the miR-17∼92 cluster in lung epithelial cells, resulting in failure to properly sacculate and partially penetrant perinatal lethality (Wang et al., 2016). *Hdac3* mutants also have increased expression of members of the *Mirg* locus, although a role for this group of miRNAs has not been assigned (Wang et al., 2016). Another study reports that expression of the miR-17∼92 cluster decreases over the course of embryonic development, and that transgenic overexpression of this cluster in lungs as they begin to develop causes a similar defect (Lu et. al, 2007). These findings, combined with our observation that levels of some members of the miR-17∼92 cluster (and its paralogous clusters), as well of the *Mirg* cluster, significantly increase in *Zswim8*^−/−^ lungs, suggested that the sacculation defect observed upon ZSWIM8 loss might be caused by the increased levels of these ZSWIM8-sensitive miRNAs.

One method of testing this hypothesis is to examine whether genetically offsetting the accumulation of these families can rescue the *Zswim8*^−/−^ phenotype. This type of rescue experiment has implicated the targeted degradation of the miR-3 family in contributing to the lethality observed upon losing TDMD in flies (Kingston et al., 2022), and it has implicated increased expression of the miR-17∼92 cluster in contributing to the sacculation phenotype of *Hdac3* mutants (Wang et al., 2016). However, we have not observed rescue of the *Zswim8*^−/−^ sacculation or embryonic growth phenotypes in genetic backgrounds designed to reduce the levels of key ZSWIM8-sensitive miRNAs, including miR-17,18,92^+/–^, miR-106b∼25^−/−^, maternal *Mirg*^−^, and miR-322∼351^−/−^ (Han et al., 2015; Ventura et al., 2008; Marty et al., 2016; Llobet-Navas et al., 2014). Although we cannot rule out the possibility that the sacculation defect stemmed from loss of a non-TDMD ZSWIM8 function, we suspect that this failure to rescue was instead because we did not reduce the relevant combination of miRNAs to the levels required to offset the effects of ZSWIM8 loss. The molecular basis of the sacculation defect can be revisited once the trigger sites that direct miRNA degradation are identified and mutated, as has been done in a few cases in mouse, zebrafish, and Drosophila (Bitetti et al., 2018; Kleaveland et al., 2018, Kingston et al., 2022). Such experiments can potentially elevate ZSWIM8-sensitive miRNAs to the status of validated TDMD substrates and open new opportunities for exploring the biological roles of this pathway.

### Sensitivities of miRNAs to ZSWIM8

We found 51 miRNAs, representing 43 miRNA families, to be ZSWIM8-sensitive in at least one of the 12 embryonic tissues examined. These 51 included 22 that had previously been identified in MEFs or iNeurons (Shi et al., 2020), but not seven others that were only identified in the cell lines. They also did not include miR-30b/c or miR-221/222, which had previously been reported to be TDMD substrates in 3T9 mouse fibroblasts and three human cell lines, respectively (Ghini et al., 2018; Li et al., 2021). Perhaps more sensitive approaches focusing on purer cell populations, or examination of other life stages, will unearth evidence that these miRNAs are indeed TDMD substrates in the animal.

ZSWIM8 sensitivity typically varied, depending on cell or tissue context (Fig. 4A–B), suggesting that TDMD selectively sculpts miRNA levels in different contexts to enhance the complexity of miRNA expression landscapes. This variable sensitivity presumably reflected the expression profiles of the trigger species, in that more effective triggers tend to be highly expressed, and TDMD efficacy can correlate with trigger expression (de la Mata et al., 2015; Shi et al., 2020, Kingston et al., 2022). A related factor is the relative stoichiometry between the miRNA and its trigger. In a study of artificial TDMD in cell culture, miRNA fold changes decrease as miRNA production increases (de la Mata et al., 2015). In tissues, the sensitivities to TDMD are presumably more complex—expected to reflect the respective cell-type compositions and the attending variation in levels of miRNAs, triggers, and protein components of the pathway. Nonetheless, miR-7 is least sensitive to Cyrano loss in the tissues in which miR-7 is most highly expressed (pituitary and pancreatic islets) (Kleaveland et al., 2018), suggesting that trigger saturation might yield diminishing fold changes in tissues in which miRNA levels are high.

Despite our use of miRNA fold change when reporting ZSWIM8 sensitivity, this metric does not tell the whole story. Increasing production of a miRNA undergoing TDMD can lower the fold change of the miRNA but increase the number of miRNA molecules that are degraded (de la Mata et al., 2015). Thus, miRNAs that undergo the greatest fold changes upon loss of TDMD might not be the ones with the most vigorous or the most biologically consequential turnover. Indeed, when analyzing the effects of ZSWIM8 loss on predicted miRNA targets, we found that the absolute change in miRNA level was more consequential than the fold change.

Many questions abound concerning the endogenous regulatory functions of ZSWIM8-mediated control of miRNA stabilities through TDMD. Our work provides evidence for possible roles for this pathway in decoupling the ultimate expression of certain miRNAs from that of other RNAs with which they share common production. For example, by our estimates, 280 miRNA hairpins – nearly half of all miRNAs encoded in the mouse genome – are produced from 60 clustered loci whose members are typically co-transcribed, presenting a possible need for post-transcriptional modulation of their levels. We found that ZSWIM8-sensitive miRNAs are disproportionately produced from genomic clusters (Fig. 6B), hinting at a role for TDMD in shaping the expression of cluster members by decoupling their accumulation in a cell-type-specific manner, dependent on the differential expression of triggers.

An extreme example of this can be found in *Mirg*, which contains the largest miRNA cluster in the mammalian genome, and is part of the imprinted, maternally expressed *Dlk1-Dio3* region on Chromosome 12 (Fig. 6A). *Mirg* miRNAs are thought to antagonize the paternal expression program (Whipple et al., 2020), and *Mirg* knockout phenotypes (observed when the mutant allele is maternally inherited) include partially penetrant neonatal lethality, due in part to metabolic defects, as well as anxiety-related behavior in adulthood (Labialle et al., 2014; Marty et al. 2016). We identified twelve ZSWIM8-sensitive miRNAs produced from *Mirg*, and an additional five miRNAs from elsewhere in the greater imprinted region—all but one of which is maternally expressed (Fig. 6A). The directed degradation of these clustered miRNAs, while serving a decoupling function, might also have evolved as a means to favor paternal interests in the parental genomic conflict that drives mammalian genomic imprinting. Future identification of triggers for these miRNAs would aid in evaluating this model; for example, paternal expression of triggers would provide further evidence of the deployment of TDMD in parental conflict.

Another example of decoupling mediated by ZSWIM8 is that of separating the expression of the two strands produced together from the same miRNA hairpin. After the miRNA duplex is excised from the hairpin, one strand associates with AGO, whereas the other is degraded. Although preferences in AGO association play an important role in setting the ultimate balance of the strands (Khvorova et al., 2003; Schwarz et al., 2003), our results show that ZSWIM8 can push this balance in either direction—in most cases, restraining accumulation of strands apparently favored to associate with AGO, but in some cases (nine of 58 murine ZSWIM8-sensitive miRNAs), further reducing strands annotated as passengers (Table S1). This latter mode might reduce unwanted activity of passenger strands in situations in which the asymmetry of AGO association is suboptimal. In other instances, both strands arising from a duplex are annotated as guides, each with a distinct cohort of targets; here, analogous to the case of clustered miRNAs, the obligate co-production of two functional miRNAs might present another need for strand-specific regulation in order to accommodate varied cellular contexts within an organism—some of which might be advantaged by the presence of both strands, and others which might favor decoupled, asymmetric accumulation. Accordingly, we identified 14 ZSWIM8-sensitive miRNAs of this variety (Table S1), including five cases in which ZSWIM8 activity caused arm switching (Fig. 5A)

Because our study focused on late embryonic development, it did not address the question of whether mammals undergo early ZSWIM8-dependent clearance of early-embryonic miRNAs, as occurs in Drosophila and *C. elegans* (Shi et al., 2020; Donnelly et al., 2022; Kingston et al., 2022). Because mammalian embryogenesis involves a large increase in fetal mass, such clearing might not be required. Future studies on the impact of ZSWIM8 during windows of developmental time could shed light on this, and temporal resolution might provide information on the biological processes affected by the degradation of particular miRNAs, including those identified in this study. Ultimately, a detailed appraisal of the biological processes regulated by endogenous TDMD will depend on genetic experiments perturbing the RNAs presumed to direct the degradation of ZSWIM8-sensitive miRNAs – most of which remain to be discovered. Identification of these putative endogenous trigger RNAs might also improve understanding of the sequence features—both of the binding sites themselves, and also perhaps of more distal elements—that modulate TDMD efficacy and enable improved prediction and design of potent triggers. Our work expanding the known set of ZSWIM8-sensitive miRNAs and the anatomical scope of ZSWIM8 activity provides a foundation for these future efforts.

## Methods

### Mouse husbandry

Mice were group-housed in a 12 hr light/dark cycle (light between 07:00 and 19:00) in a temperature-controlled room (21.1 ± 1.1°C) at the Whitehead Institute for Biomedical Research with free access to water and food and maintained according to protocols approved by the Massachusetts Institute of Technology Committee on Animal Care. Euthanasia of adults was performed by CO_2_ inhalation. Sex was not determined for embryos or neonatal pups, except where indicated. Embryos were weighed after being patted dry with a paper towel.

### Generation of mutant mice

Mice with mutations in *Zswim8* were generated by injecting injecting one-cell C57BL/6J embryos with Cas9 protein complexed with a sgRNA designed to cut within exon 2 of *Zswim8* (Fig. 1A). F0 mice containing resulting deletions (1, 4, and 7 nt, respectively) and insertions (2 nt) were bred to C57BL/6J mice, and then F1 mice were crossed to generate lines with the desired heterozygous mutations (*Zswim8*^+/–^). *Zswim8*^+/–^ lines were maintained by breeding to C57BL/6J, and resulting heterozygous offspring were intercrossed to generate *Zswim8*^+/+^, *Zswim8*^+/–^, and *Zswim8*^−/−^ embryos and neonates used in this study. No substantial phenotypic differences were observed between mice bearing each of four *Zswim8* alleles, which were used interchangeably in this study.

### Genotyping

Genomic DNA was extracted from mouse ear punches using the HotSHOT method (Truett et al., 2000). For *Zswim8* mutants, PCR was performed with KAPA HiFi HotStart ReadyMix (Roche), and amplicons were purified (QiaQuick PCR Purification Kit, QIAGEN) and submitted for Sanger sequencing. Primers, PCR conditions, and expected amplicon sizes are listed in Table S1.

### RNA extraction

For mouse tissue, total RNA was extracted with TRI Reagent according to the manufacturer’s protocol with the following modifications. Tissues from E18.5 embryos were rapidly dissected after euthanasia of pregnant dams (CO_2_) and flash-frozen using liquid nitrogen. Tissue was transferred to a 50 mL conical tube, 1–2 mL of TRI Reagent was added depending on the tissue volume, and the tissue was homogenized with a TissueRuptor (QIAGEN) and disposable probes. Phase separation was performed by adding 100 μL 1-Bromo-3-chloropropane (Sigma) to 1 mL of homogenate. Aqueous phase was extracted and precipitated with isopropanol, and the pellet was washed twice with 75% ethanol. All RNA was resuspended in RNAse-free water.

### Bulk RNA-seq

Bulk RNA-seq data were generated from total RNA isolated from tissues dissected from one E18.5 *Zswim8*^+/–^ embryo and one *Zswim8*^−/−^ embryo selected from the same litter. This procedure was repeated with another litter, for *n* = 2 biological replicates. RNA-seq libraries were prepared from 1 μg samples of total RNA using the KAPA RNA HyperPrep Kit with RiboErase (HMR) kit (KAPA Biosystems), and sequenced on the Illumina NovaSeq platform with 50-nt paired-end reads. Reads were aligned to the mouse genome (mm10) using STAR v2.7.1a (Dobin et al., 2013) with the parameters ‘‘–runThreadN 24 –outFilterMultimapNmax 1 – outFilterMismatchNoverLmax 0.04 –outFilterIntronMotifs RemoveNoncanonicalUnannotated – outSJfilterReads Unique –outReadsUnmapped Fastx –quantMode GeneCounts –outSAMtype BAM SortedByCoordinate”. Aligned reads were assigned to genes using annotations from RefSeq (downloaded August 3, 2022) and counted using htseq-count v0.6.1 (Anders et al., 2015). Further analyses were performed on genes passing the expression threshold of at least five counts in each of the four libraries from a tissue. Counts arising from *Xist* or *Tsix* were used to assign sex (one *Zswim8*^+/–^ embryo was female; the other three embryos were male), and removed before proceeding with analysis. No substantial differences in ZSWIM8 sensitivity were observed between tissues of male and female embryos (Fig. S3F). Transcripts per million (TPM) values were computed by dividing the counts mapping to each gene by the length of its longest annotated mRNA isoform, and then dividing by the sum of this measure across all genes, and multiplying by 10^6^. Differential expression and significance levels were determined using DESeq2 v1.26.0 (Love et al., 2014) without the lfcShrink() function.

### sRNA-seq

sRNA-seq data were generated from the same total-RNA samples subjected to bulk RNA-seq, as described in the preceding section. Libraries were prepared and sequenced from 5 µg of total RNA as described for mammalian samples (Shi et al., 2020). Synthetic miRNA spike-in RNA oligonucleotides were added to each total-RNA sample prior to size-selection, in proportion to measured RNA content. A detailed protocol for constructing sRNA-seq libraries is available at http://bartellab.wi.mit.edu/protocols.html. Libraries were sequenced on the Illumina NovaSeq platform with 100-nt single-end reads. Processing of sequencing read data and subsequent analyses was as described (Shi et al, 2020). Counting of annotated miRNAs was performed by string-matching the first 19 nt of each read to a dictionary derived from the set of mature miRNA names and sequences downloaded from TargetScan7 Mouse (Agarwal et al., 2015). These dictionaries were primarily generated from the mature miRNA sequences whose first 19 nt were unique, to avoid ambiguities caused by 3′ alterations, as described (Shi et al., 2020). For the 48 annotated miRNAs whose first 19 nt were not unique, all miRNAs sharing a 19-nt prefix were collapsed to create a single dictionary entry with sequence of the 19-nt prefix, and given a name that combined the names of the collapsed miRNAs (e.g., mmu-miR-466j/mmu-miR-466m-5p/mmu-miR-669m-5p). This procedure added an additional 21 entries to the dictionary.

Further analyses were performed on miRNA or passenger-strand species passing the expression threshold of at least five matching reads in each of the four libraries derived from a tissue. When calculating normalized abundances, the mean of miRNA counts across two replicates was taken, normalized to the sum of all mean miRNA-matching counts (after removing counts accruing to the synthetic spike-in oligos), and multiplied by 10^6^ to yield counts per million (CPM). Differential expression and significance levels were determined using DESeq2 v1.26.0 (Love et al., 2014) without use of the lfcShrink() function, using raw counts as input. For all plotting and analyses, fold changes observed between *Zswim8*^+/–^ and *Zswim8*^−/−^ samples, as well as their standard errors, were generated by DESeq2.

For identifying miRNAs significantly upregulated between *Zswim8*^+/–^ and *Zswim8*^−/−^ samples from a tissue, we used a method based on a modified bi-beta-uniform mixture (BBUM) model (Wang and Bartel, 2022). For each tissue, all miRNAs with an FDR-adjusted *p* value < 0.05 were called as significantly upregulated. A significantly upregulated miRNA produced from a single locus was classified as a ZSWIM8-sensitive miRNA if its log_2_ fold change upon *Zswim8* loss was significantly greater than that of its passenger strand, as would be expected of substrates of TDMD under the assumption that the miRNA’s degradation rate is affected by ZSWIM8, but not its production rate, nor the production or degradation rates of the passenger. In the case of a significantly upregulated miRNA produced from more than one locus, the analogous assumptions led to classification of the miRNA as a ZSWIM8-sensitive miRNA if its log_2_ fold change upon *Zswim8* loss was significantly greater than the log_2_ of the sum of its passenger strands’ levels in *Zswim8*^−/−^ subtracted by the log_2_ of the sum of its passenger strands’ levels in *Zswim8*^+/–^. An additional simplifying assumption was that the production rates of the miRNA and passenger produced from a locus differed by the same multiplicative factor for all loci producing the miRNA. For these analyses, the aforementioned ‘significantly greater than’ condition was achieved if the two compared quantities were separated beyond their standard errors, as calculated by DESeq2 (1.26.0) (Love et al., 2014) or propagated therefrom. Any significantly upregulated miRNA that did not meet this standard of significance was classified, together with its passenger strand, as indirectly ZSWIM8-sensitive. Additional miRNAs that were not called as significantly upregulated were also classified as ZSWIM8-sensitive if they showed increases in at least 11 of 12 *Zswim8*^−/−^ tissues, with a median log_2_ fold-change > 0.2 across tissues, as well as a greater increase compared to that of its cognate strand, with a median difference in log_2_ fold-change > 0.2 across tissues. Nine miRNAs met these alternative criteria: miR-212-3p, miR-345-3p, miR-351-5p, miR-361-5p, miR-380-3p, miR-466m-3p, miR-744-3p, miR-99b-3p, and miR-497a-3p. In the few cases (e.g., 2 of 33 ZSWIM8-sensitive miRNAs in forebrain) in which the passenger strand was not detected above the count cutoff in a tissue, the significantly upregulated miRNA was classified as ZSWIM8-sensitive.

A miRNA that was not called ZSWIM8-sensitive in a tissue was considered marginally sensitive in that tissue if it was classified as ZSWIM8-sensitive in at least one other tissue, MEF, or iNeuron (Shi et al., 2020) and if its log_2_ fold change was significantly greater than that of its passenger strand in the tissue under consideration, as assessed by the method based on standard errors described above. In rare cases (miR-335-3p, miR-429-5p, miR-466i-3p, miR-544-3p, miR-652-5p, miR-764-5p, miR-497a-3p, miR-744-3p, miR-99b-3p), the annotated passenger strand was ZSWIM8-sensitive, in which case, for purposes of our analyses, the ZSWIM8-sensitive strand was considered the guide strand, and the other strand was considered the passenger strand.

From the union of miRNAs annotated as ZSWIM8-sensitive across the twelve profiled tissues, six were manually removed and recategorized as indirectly ZSWIM8-sensitive (miR-743b-3p, miR-883a-3p, miR-883b-3p, miR-881-3p, miR-880-3p, and miR-742-3p). These six were likely erroneously categorized because they are members of a miRNA cluster that was likely transcriptionally up-regulated in placenta (Fig. S3E), but unlike the other detected miRNAs in the cluster, had passenger strands that did not meet our detection threshold. In addition, two (miR-451a and miR-122-5p) were removed on the basis of possible contamination with small quantities of material from blood or liver during dissection, and two more (miR-429-3p and miR-135b-5p) were removed because they were down-regulated in many tissues except for one, and were thus possibly misannotated by our criteria by chance. miRNAs classified as ZSWIM8-sensitive were examined within individual replicate libraries to ensure that classification did not appear to be driven by sex differences between the samples (see preceding section).

For analysis of miRNA clusters, mouse miRNA annotations were from TargetScan7 (Agarwal et al., 2015) and their genomic coordinates and associated primary transcript sequences were from miRBase Release 22.1 (Kozomara et al., 2019). To focus on confidently annotated miRNAs, a miRNA hairpin locus was only considered for analysis if at least one of its strands was annotated by TargetScan7 as part of a ‘broadly conserved’, ‘conserved’, or ‘poorly conserved but confidently annotated’ miRNA family. A cluster was defined as a set of at least two miRNAs hairpin loci (annotated in miRBase as miRNA primary transcripts), on the same chromosomal strand whose boundaries fell within a contiguously sliding 10 kb window. By this definition, we found the mouse genome to contain 604 annotated miRNA loci, of which 280 collectively resided within 60 clusters. A hairpin locus was counted as encoding a ZSWIM8-sensitive miRNA if either mature strand produced from it was annotated as ZSWIM8-sensitive in an E18.5 tissue, MEF or iNeuron (Shi et al., 2020). Four ZSWIM8-sensitive miRNAs were each annotated as encoded by two separate loci (miR-92a-3p, miR-194-5p, miR-7-5p, and miR-29b-3p), and each of these eight hairpins was counted as a ZSWIM8-sensitive miRNA hairpin locus.

For arm-switching analysis, the passenger strands produced from all miRNA loci encoding a given ZSWIM8-senstive miRNA were pooled and treated identically, and this pooled passenger strand was compared in abundance to the corresponding ZSWIM8-sensitive strand between *Zswim8*^+/–^ and *Zswim8*^−/−^ tissues. An arm switching event was counted in a given tissue if the identity of the more abundant of the two strands associated with a ZSWIM8-sensitive miRNA was observed to depend on *Zswim8* genotype.

### scRNA-seq

scRNA-seq data were generated from dissociated whole lungs dissected from two *Zswim8*^+/–^ and three *Zswim8*^−/−^ E18.5 embryos. For dissociation, each set of lungs was minced on ice with scissors and incubated in 500 μL digestion solution composed of 100 μL collagenase (2000U/mL; Thermo Fisher Scientific), 16.5 uL DNAse I (0.33U/mL; Roche), and 383.5 μL DMEM/F-12 media (Life Technologies), and rotated end-over-end for 45 minutes at 37°C. Following dissociation, 500 μL DMEM/F-12 + 10% fetal bovine serum (FBS) was added to the reaction, and the suspension was filtered through a 70 μm strainer. The filtrate was pelleted at 400*g* for 5 min at 4°C, resuspended in 500 μL red blood cell lysis buffer (StemCell Technologies) with a wide-bore P1000 tip, incubated for one minute at room temperature, and again pelleted at 400*g* for 5 min at 4°C. The pellet was resuspended in 200 μL DMEM/F12 + 10% FBS with a wide-bore P1000 tip, and cells were counted and assessed for viability by trypan blue staining. For the five samples, viabilities ranged from 64–81%. Cells were then loaded onto a Chromium controller for library preparation using the Chromium Next GEM Single Cell 3′ GEM, Library & Gel Bead Kit v3.1 (10X Genomics), targeting 10,000 cells per library, and libraries were sequenced on an Illumina NovaSeq platform with 150-nt paired-end reads.

Sequencing data was aligned to the mouse genome (mm10) and demultiplexing, barcode processing, gene counting and aggregation were performed using the Cell Ranger software v4.0.0 (https://support.10xgenomics.com/single-cell-gene-expression/software/pipelines/latest/what-is-cell-ranger). Processed data was further filtered and analyzed using ScanPy (Wolf et al., 2018). Each library was filtered for cells with detectable counts for >200 and <12,000 genes, each detected in at least 3 cells, a mitochondrial gene-count percentage of <15%, and a hemoglobin gene-count percentage of <10%. Contributions per cell from total gene counts, percent mitochondrial gene counts, and percent hemoglobin gene counts were then regressed out. For analyses of all lung cells, the data from all libraries were combined and UMAP was performed with 10 nearest neighbors and 40 principal components, and neighborhood detection was performed using the Leiden clustering method, using a resolution parameter of 0.2. For analyses of alveolar epithelial cells, the cluster characterized by expression of the *Nkx2-1* lineage marker was extracted, as described in the text. On this subset, UMAP was performed with five nearest neighbors and 15 principal components, and clustering was performed with a resolution parameter of 0.4.

### miRNA targeting analysis

Predictions of miRNA targets were from TargetScan7 Mouse (Agarwal et al., 2015). Using these annotations, the set all predicted targets of the miRNA family was taken as the least stringent cohort used for analyses. The set of all conserved predicted targets was taken as the cohort with medium stringency, and the subset of all predicted targets in the top decile of cumulative weighted context++ scores, named the top predicted targets, was taken as the cohort with highest prediction stringency. Finally, the set of all transcripts represented, excluding those predicted as targets of a given family, were used as the pool of transcripts not predicted to be targeted by that family, from which the no-site cohorts were selected.

For analyses of miRNA-mediated repression in a given tissue, the cohorts described above were filtered for transcripts with expression >10 TPM in that tissue for all datasets from both *Zswim8*^+/–^ and *Zswim8*^−/−^ genotypes. For each cohort of predicted targets, a matched no-site cohort was sampled, under the condition that for each predicted target, five no-site transcripts were selected from a bin with matched 3′-UTR length. The 3′-UTR bins were generated by partitioning by length all 3′ UTRs into 20 uniformly populated bins, ranging from the shortest to longest annotated 3′ UTRs in TargetScan.

To compute repression of a predicted-target cohort in a tissue, the distribution of log_2_ fold changes in *Zswim8*^−/−^ relative to *Zswim8*^+/–^ was compared to that of a corresponding no-site cohort using the Mann-Whitney U Test, and the median log_2_ fold change of the predicted-target cohort was subtracted by that of the no-site cohort to yield a measure of fold repression attributable to a miRNA family. This procedure was repeated 21 times, each with a different randomly sampled no-site cohort, to yield a mean and standard error for fold repression. These values, along with the median *p* value from the 21 samplings (Mann-Whitney U Test), are reported for each of the predicted-target cohorts for each of the three most affected miRNA families in each tissue (Fig. 6).

### Histology and immunofluorescence microscopy

For general histological analysis of heart and lungs, E18.5 embryos were dissected while submerged under ice-cold PBS to prevent breathing of air and preserve developmental anatomy, and the thoracic cavity was exposed for overnight fixation with Bouin’s solution, followed by paraffin embedding, serial microtome sectioning with a thickness of 5 µm, and staining with hematoxylin and eosin (H&E). For immunofluorescence experiments, lungs dissected as described above were fixed in 4% paraformaldehyde (Electron Microscopy Sciences) overnight at 4°C, transferred to 30% sucrose for cryoprotection, embedded in Tissue-Tek optimal cutting temperature compound (Sakura Finetek), and sectioned at a thickness of 25 µm using a cryostat microtome (Leica). Tissue sections were stained overnight with the following primary antibodies: hamster anti-Podoplanin (8.1.1, DHSB; 1:800 dilution), rabbit anti-pro-Surfactant C (AB3786, Millipore; 1:1500 dilution), and for one hour at room temperature with the following fluorescent secondary antibodies: goat anti-hamster IgG AlexaFluor 568 (Invitrogen; 1:500 dilution), goat anti-rabbit IgG AlexaFluor 647 (Invitrogen; 1:500 dilution), as well as with NucBlue Hoechst 33342 (Invitrogen; two drops/mL). Fluroescence microscopy was performed on a Nikon Ti widefield microscope.

## Data access

All raw and processed sequencing data generated in this study have been submitted to the NCBI Gene Expression Omnibus (GEO; https://www.ncbi.nlm.nih.gov/geo/) under accession number GSE231450.

## Acknowledgments

We thank M. Frank for assistance with tissue dissections; R. Bronson for assistance with histological analysis; D. Lin, M. Frank, R. Saunders, L. Blodgett, A. Latifkar, E. Kingston, S. McGeary, V. Auyeung, and J. Rajagopal for helpful discussions; A. Ventura, J. Silva, and J. Cavaille for sharing mouse strains; the Whitehead Genetically Engineered Models Center for assistance with generating mouse strains; the Whitehead Genome Technology Core for sequencing; the Koch Institute’s Swanson Biotechnology Center’s Histology Core for histology; the W.M. Keck Microscopy Facility for imaging.

## Author contributions

C.Y.S. and D.P.B. conceived the project and designed the study. C.Y.S. and J.S. performed mouse husbandry and tissue processing, with assistance from B.K. L.E.E. and C.Y.S. prepared samples for single-cell RNA-sequencing. C.Y.S. prepared all other sequencing libraries and performed sequencing data analysis. C.Y.S. and R.R.C. performed immunofluorescence microscopy. B.K. and R.R.C. advised on pathological analysis. C.Y.S. and D.P.B. drafted the manuscript.

