## Supplemental Figures for "The widespread influence of ZSWIM8 on microRNAs during mouse embryonic development"

### Supplementary Information

#### Table S1: ZSWIM8-sensitive miRNAs across tissues

### Supplemental Figures and Legends

Shi_FigS1

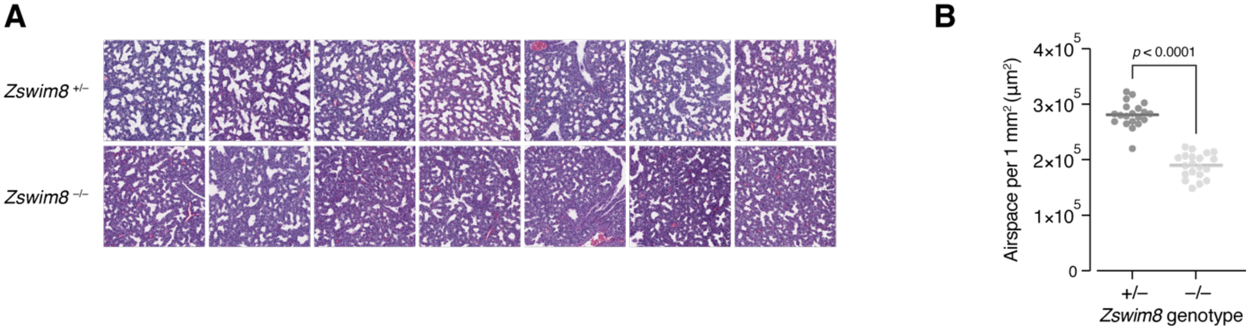


**Figure S1.** Airspace defect in *Zswim8*$^{-/-}$ embryonic lung; related to Figure 2. (A) Images of representative H&E-stained fixed sections from lungs of a E18.5 *Zswim8*$^{+/-}$ embryo and a *Zswim8*$^{-/-}$ littermate. (B) Quantification of airspace area from 20 sections with adjacent sections spaced >200 µm apart. *p*-value from Mann-Whitney test.

Shi_FigS2

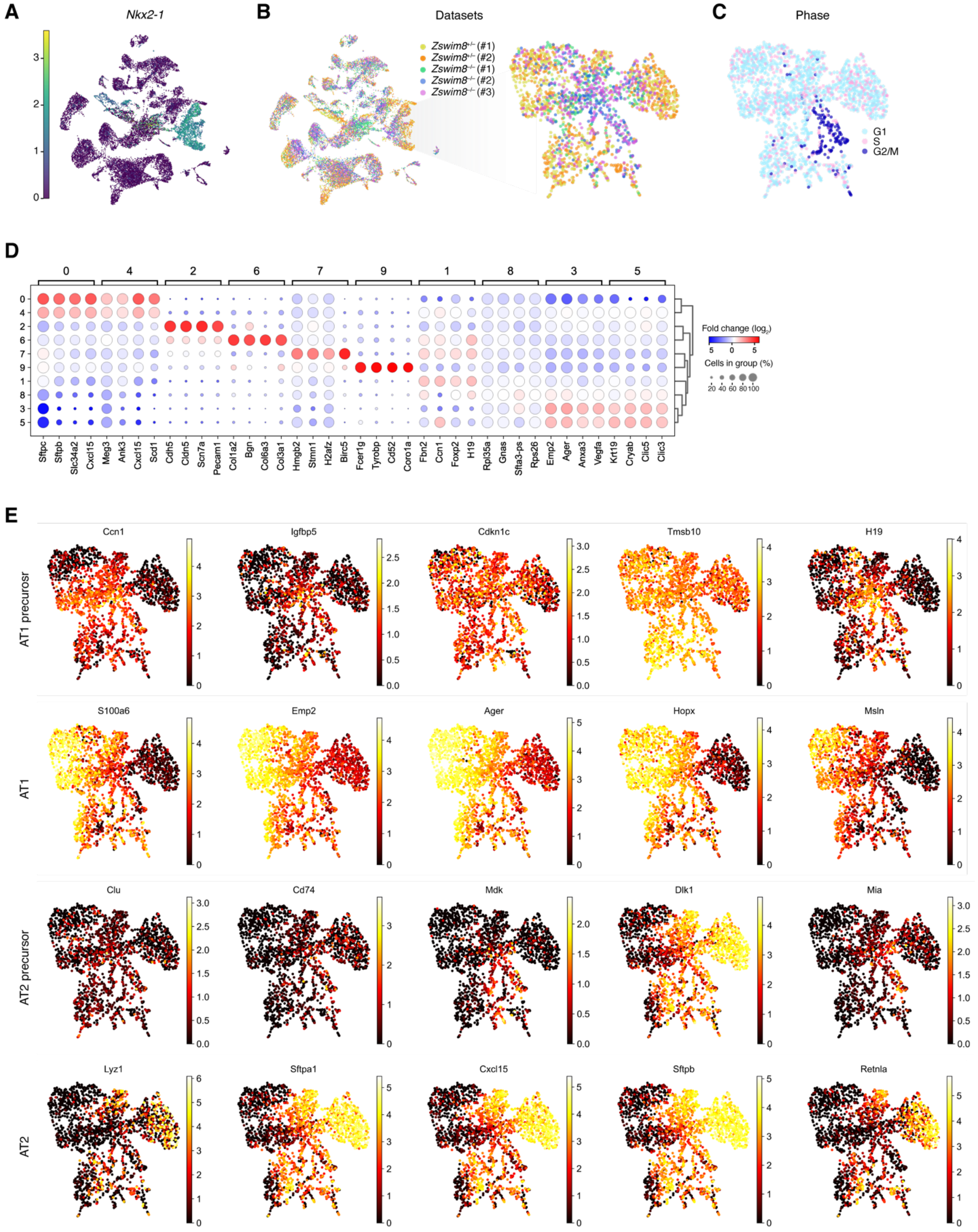
A
Nkx2-1
B
Datasets
Zswim8+/- (#1)
Zswim8+/- (#2)
Zswim8-/- (#1)
Zswim8-/- (#2)
Zswim8-/- (#3)
C
Phase
G1
S
G2/M
D
Fold change (log2)
Cells in group (%)
Sftpc Sftpb Slc34a2 Cxcl15 Meg3 Ank3 Cxcl15 Scd1 Cdh5 Cldn5 Scn7a Pecam1 Col1a2 Bgn Col6a3 Col3a1 Hmgb2 Stmn1 H2afz Birc5 Fcer1g Tyrobp Cd52 Coro1a Fbn2 Ccn1 Foxp2 H19 Rpl35a Gnas Sfta3-ps Rps26 Emp2 Ager Anxa3 Vegfa Krt19 Cryab Clic5 Clic3
E
AT1 precurosr
Ccn1
Igfbp5
Cdkn1c
Tmsb10
H19
AT1
S100a6
Emp2
Ager
Hopx
Msln
AT2 precursor
Clu
Cd74
Mdk
Dlk1
Mia
AT2
Lyz1
Sftpa1
Cxcl15
Sftpb
Retnla

**Figure S2.** Single-cell RNA-sequencing from lungs of *Zswim8*$^{-/-}$ embryos; related to Figure 3. (A) The three most significantly enriched genes in each unsupervised cluster generated from all cells captured; related to Fig. 3A. Row labels indicate cluster number, and column labels indicate enriched genes grouped by cluster number. Rows are hierarchically clustered. Size of dots indicates the fraction of cells in given cluster with detectable counts for a given gene. Fill color of dots indicates the fold difference in mean expression in a given cluster, relative to the union of all other clusters; color bar is in units of $\log_2$. Significance determined by Wilcoxon rank-sum test, adjusted by the Benjamini-Hochberg method. (B) UMAP of all cells captured, as in Fig. 3A, with color indicating expression of the canonical lung epithelial marker *Nkx2-1*. Color bar is in units of log(1+ CP10K); related to Fig. 3A. (C) Left: As in Fig. 3A, UMAP of all cells captured, with color indicating individual dataset; related to Fig. 3A. Right: As in Fig. 3B, UMAP re-embedding of cells of Cluster 2 shown in Fig. 3A (corresponding largely to *Nkx2-1*-positive cells), with color indicating individual dataset; related to Fig. 3B. (D) As in Fig. 3B, UMAP re-embedding of cells of Cluster 2 shown in Fig. 3A (corresponding largely to *Nkx2-1*-positive cells), with color indicating inferred cell cycle phase; related to Fig. 3B. (E) The four most significantly enriched genes in each cluster generated from re-clustering of re-embedded of cells of Cluster 2 shown in Fig. 3A; related to Fig. 3B. (F) Expression of reported marker genes for AT1 precursor, AT1, AT2 precursor, and AT2 cells (Frank et al., 2019) in re-embedding of cells of Cluster 2 shown in Fig. 3A; related to Fig. 3B–C.

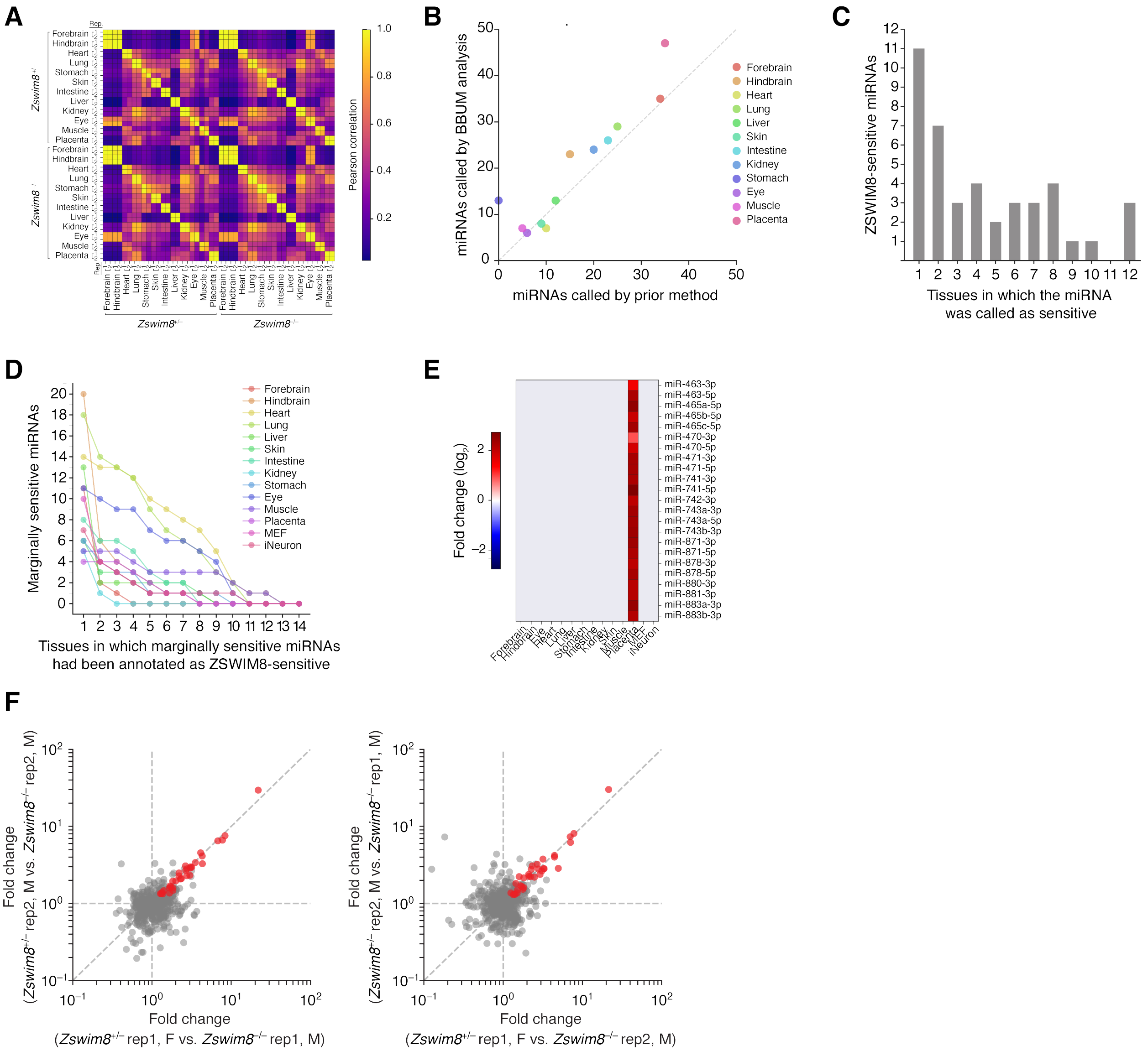


**Figure S3.** Influence of ZSWIM8 on miRNAs across embryonic tissues; related to Figure 4. (A) Correspondence of miRNA levels between E18.5 embryonic tissues of two biological replicates, as measured by sRNA-seq. Color map indicates Pearson correlation. (B) Number of significantly ZSWIM8-sensitive miRNAs in E18.5 embryonic tissues called by BBUM analysis (Wang et al., 2022), compared to that called by our original method (Shi et al., 2020). (C) The number of miRNAs called as ZSWIM8-sensitive in E18.5 embryos by both significant upregulation in BBUM analysis and significant elevation of guide strand above passenger strand, compared to the number of tissues in which they were independently called as ZSWIM8-sensitive. (D) The frequency by which marginally sensitive miRNAs were independently called as ZSWIM8-sensitive in other tissues. Shown for each tissue are the number of marginally sensitive miRNAs plotted as a function of the number of other datasets (embryonic tissues, as well as MEF and iNeuron cells (Shi et al., 2020)) in which these miRNAs were independently called as ZSWIM8-

sensitive. (E) Changes in levels of all detected members of an X-linked genomic cluster encoding miR-743a, miR-743b, miR-742, miR-883a, miR-883b, miR-471, miR-741, miR-463, miR-880, miR-878, miR-881, miR-871, miR-470, miR-465d, miR-465c, miR-465b, and miR-465a. Color bar indicates $\log_2$ fold change; gray, not detected. (F) Correspondence of ZSWIM8 sensitivity of miRNAs in forebrain between individual samples derived from male (M) and female (F) embryos of two biological replicates, as measured by sRNA-seq. Shown are comparisons of fold changes in miRNA levels between samples of the indicated sexes and genotypes. miRNAs called as ZSWIM8-sensitive in forebrain are shown in red. Similar results (not shown) were observed in the eleven other embryonic tissues.

Shi_FigS4

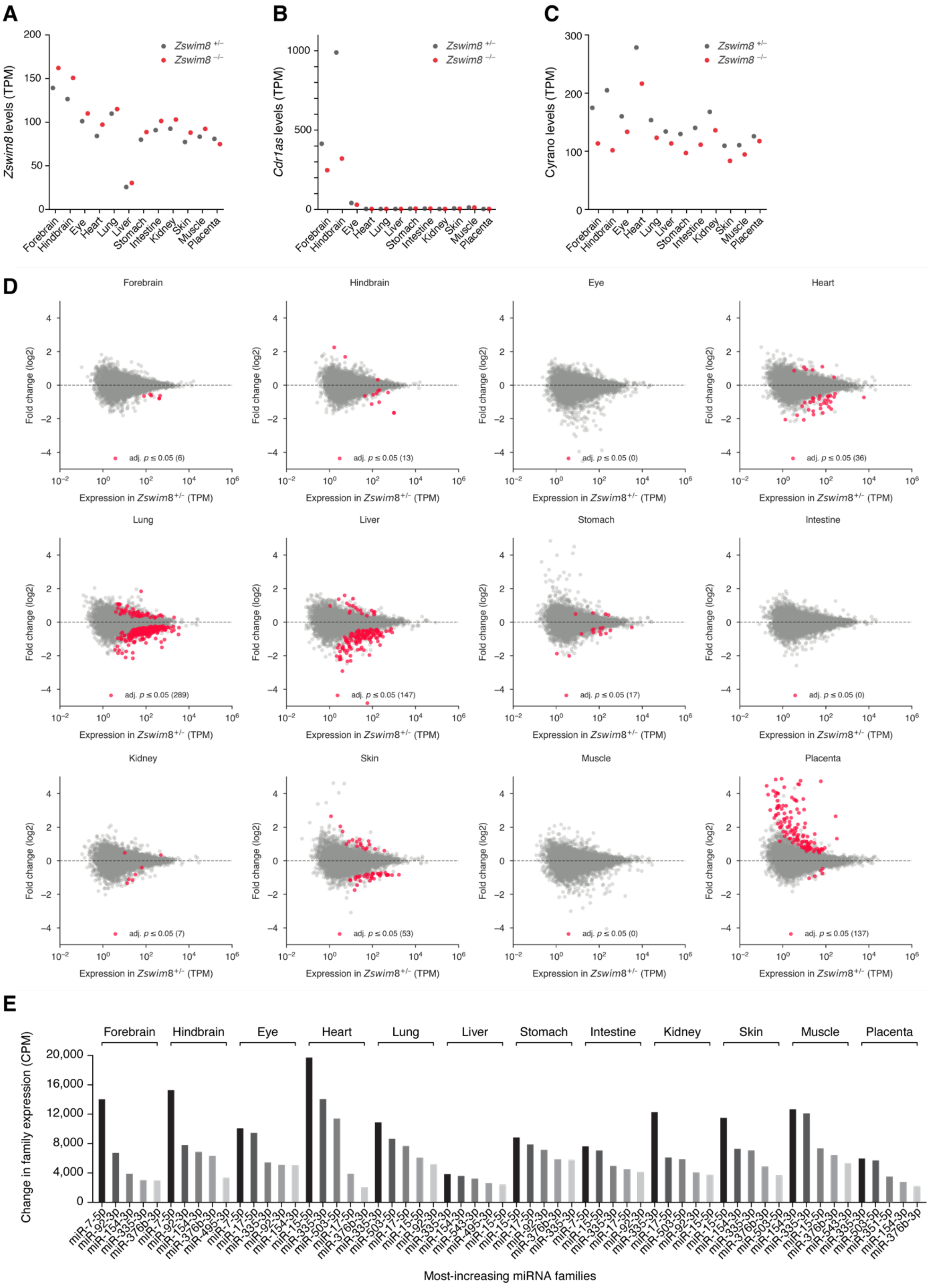
A
Zswim8 levels (TPM)
Zswim8 +/−
Zswim8 −/−
B
Cdr1as levels (TPM)
C
Cyrano levels (TPM)
Forebrain
Hindbrain
Eye
Heart
Lung
Liver
Stomach
Intestine
Kidney
Skin
Muscle
Placenta
D
Fold change (log2)
Expression in Zswim8+/- (TPM)
Forebrain adj. p ≤ 0.05 (6)
Hindbrain adj. p ≤ 0.05 (13)
Eye adj. p ≤ 0.05 (0)
Heart adj. p ≤ 0.05 (36)
Lung adj. p ≤ 0.05 (289)
Liver adj. p ≤ 0.05 (147)
Stomach adj. p ≤ 0.05 (17)
Intestine adj. p ≤ 0.05 (0)
Kidney adj. p ≤ 0.05 (7)
Skin adj. p ≤ 0.05 (53)
Muscle adj. p ≤ 0.05 (0)
Placenta adj. p ≤ 0.05 (137)
E
Change in family expression (CPM)
Forebrain: miR-7-5p, miR-92-3p, miR-154-3p, miR-335-3p, miR-376b-3p
Hindbrain: miR-7-5p, miR-92-3p, miR-154-3p, miR-376b-3p, miR-495-3p
Eye: miR-7-5p, miR-17-5p, miR-335-3p, miR-92-3p, miR-154-3p
Heart: miR-15-5p, miR-335-3p, miR-503-5p, miR-17-5p, miR-376b-3p
Lung: miR-335-3p, miR-503-5p, miR-17-5p, miR-15-5p, miR-92-3p
Liver: miR-335-3p, miR-154-3p, miR-543-3p, miR-495-3p, miR-15-5p
Stomach: miR-15-5p, miR-17-5p, miR-92-3p, miR-376b-3p, miR-335-3p
Intestine: miR-7-5p, miR-15-5p, miR-335-3p, miR-17-5p, miR-92-3p
Kidney: miR-335-3p, miR-17-5p, miR-503-5p, miR-92-3p, miR-15-5p
Skin: miR-15-5p, miR-154-3p, miR-335-3p, miR-376b-3p, miR-503-5p
Muscle: miR-154-3p, miR-335-3p, miR-15-5p, miR-376b-3p, miR-543-3p
Placenta: miR-335-3p, miR-503-5p, miR-351-5p, miR-154-3p, miR-376b-3p
Most-increasing miRNA families

**Figure S4.** Influence of ZSWIM8 on levels of miRNA targets across embryonic tissues; related to Figure 7. (A) Expression of *Zswim8* tissues of *Zswim8*$^{+/–}$ and *Zswim8*$^{–/–}$ E18.5 embryos across two biological replicates, as quantified by RNA-seq. TPM; transcripts per million. (B) Expression of the circular RNA *Cdr1as* in tissues of *Zswim8*$^{+/–}$ and *Zswim8*$^{–/–}$ E18.5 embryos. (C) Expression of the long noncoding RNA Cyrano in tissues of *Zswim8*$^{+/–}$ and *Zswim8*$^{–/–}$ E18.5 embryos. (D) Expression levels of RNA species in tissues of E18.5 embryos, as quantified by RNA-seq. Shown are fold changes in levels of *Zswim8*$^{–/–}$ tissues, relative to *Zswim8*$^{+/–}$. Highlighted are differentially expressed species with adjusted $p$-value < 0.05 as computed by DESeq2 (Love et al., 2014). (E) Changes in the levels of miRNA seed-family members, as quantified by sRNA-seq. Shown are the five most-increasing families, as defined by total changes in depth-normalized counts per million (CPM), in *Zswim8*$^{–/–}$ E18.5 tissues, relative to *Zswim8*$^{+/–}$; related to Fig. 7B.
